# The *Trypanosoma cruzi* cell atlas; a single-cell resource for understanding parasite population heterogeneity and differentiation

**DOI:** 10.1101/2024.10.01.616042

**Authors:** Ross Laidlaw, Marta Garcia Sanchez, Juliana Da Silva Pacheco, Luciana De Sousa Paradela, Thomas D Otto, Manu De Rycker

## Abstract

*Trypanosoma cruzi*, the causative agent of Chagas disease, exhibits a complex life cycle with multiple hosts, stages and differentiation steps. We present a complete cell atlas for the *T. cruzi* life cycle, based on single cell transcriptomes for over 31,000 cells and population-based transcriptomics. The atlas reveals many life cycle associated genes and can be utilised to accurately annotate life cycle stages. It provides detailed insights into cell heterogeneity, including cell-specific repertoires of surface antigens in trypomastigotes, with key implications for immune responses. Enabled by single-cell resolution, we define the transcriptomic changes that occur across the epimastigote to metacyclic trypomastigote differentiation axis. Furthermore, we provide comprehensive UTR annotation, identifying previously unannotated transcripts as well as revealing alternative poly-adenylation and an unanticipated complexity of reverse strand and antisense transcripts. This *T. cruzi* atlas provides a comprehensive resource and unlocks a range of new avenues for research on this important human pathogen.

## Main

*Trypanosoma cruzi*, a protozoan parasite of the kinetoplastida class, is the aetiologic agent of Chagas disease, a neglected tropical disease and lifelong debilitating zoonosis that affects approximately 6–8 million people worldwide^1^. Current therapeutic options are limited to benznidazole and nifurtimox, with important side effects and limited efficacy in the chronic stage of the disease. *T. cruzi* exhibits a complex life cycle with at least four distinct stages in two hosts, an invertebrate triatomine vector and a broad range of mammalian species, including humans. Motile metacyclic trypomastigotes, present in the faeces of the vector, infect mammalian hosts through the insect bite wound, damaged skin or mucosal surfaces and invade host cells^2,3^. Differentiation into non-motile, replicative, amastigotes starts in parasitophorous vacuoles, followed by escape to the host cell cytosol, where differentiation is completed^4^. After multiple cycles of division, amastigotes differentiate into non-replicative trypomastigotes which, upon cell rupture, are released into the bloodstream, facilitating dissemination. Blood trypomastigotes ingested by the vector migrate along the insect digestive tract and differentiate into replicative epimastigotes when reaching the midgut. Once in the rectal ampulla, nutritional stress induces metacyclogenesis, with epimastigotes differentiating into non-replicative metacyclic trypomastigotes, which are released in the faeces of the bug, closing the cycle^2,3,5^. This complex life cycle necessitates precise regulation of gene expression. Common to other trypanosomatids, transcription initiation in *T. cruzi* is not regulated at the level of individual genes. Transcription is polycistronic, followed by a trans-splicing process and polyadenylation to generate individual mRNAs. Gene expression is regulated through multiple mechanisms at the levels of mRNA processing, translation and degradation^6^.

Transcriptomics studies have shown substantial changes in mRNA levels as the parasite progresses through its life cycle^7,8^, including in genes with known stage-related biological functions, such as host cell invasion or metabolic adaptation to changing environments^9–12^. These transcriptome studies combined with other approaches^2,13–18^, have significantly advanced our understanding of the *T. cruzi* life cycle and indicate a level of complexity beyond the four canonical stages described above, including the recently described dormant or persister amastigotes, which have important relevance for the development of therapeutics^19–22^. Studying transient forms is difficult using bulk methods, as they tend to mask the heterogeneity within the tested populations. With the advent of single-cell transcriptomic methods this challenge can now be addressed, as demonstrated by studies on other parasitic protozoa such as *Plasmodium spp* ^23–28^, *Trypanosoma brucei*^29–31^, *Toxoplasma gondii*^32^ and *Babesia spp*^33^.

Here, we present a single-cell RNA-seq atlas of the *T. cruzi* life cycle as well as parallel bulk RNA-seq data. Our findings show that both techniques offer complementary insights, and that combining the approaches presents an effective strategy for unravelling the parasite’s transcriptional dynamics during stage transitions and to accurately annotate UTRs. The powerful dataset details the dynamic changes across the epimastigote to metacyclic trypomastigote differentiation axis and uncovers distinct trypomastigote sub-populations and cell-specific repertoires of surface proteins. This single-cell atlas provides a key resource for further study of the transcriptional dynamics during the *T. cruzi* life cycle.

## Results

### *Trypanosoma cruzi* life cycle cell atlas

To generate a single-cell life cycle cell atlas for *T. cruzi* we collected samples from across the life cycle (Fig. 1). Cell viability was >98% for EP, SE/MT, ADT and AMA_120h_, and > 92% for AMA_6h_ and AMA_24h_ (Supplementary Fig. 1, Supplementary Table 1). For library construction samples were pooled as indicated in Fig. 1b. The pooling strategy was determined by balancing cost and sequencing depth per life cycle stage. Pools containing no more than two life cycle stages were generated, as well as a pool of all stages. Sequencing data was mapped against the Dm28c 2018 reference genome as this provided better mapping statistics than the Sylvio X10 2017 genome (Supplementary Table 2). A large fraction of the reads mapped to rRNA genes, likely due to capture through homopolymer tracks (see methods). The median number of genes detected per cell for each sample ranged from 298 – 928 and the percentage of *T. cruzi* reads mapping confidently to the transcriptome ranged from 3.9% - 21.4% (Supplementary Table 3). After quality control, 31,065 cells were identified as valid across all the samples, with information for 15,321 genes detected in total (out of 19,112 genes in the Dm28c 2018 reference genome). Full quality control cutoffs are shown in Supplementary Table 4. After processing of the valid cells, 10 clusters were identified across the atlas (Fig. 2a).

**Figure 1.**
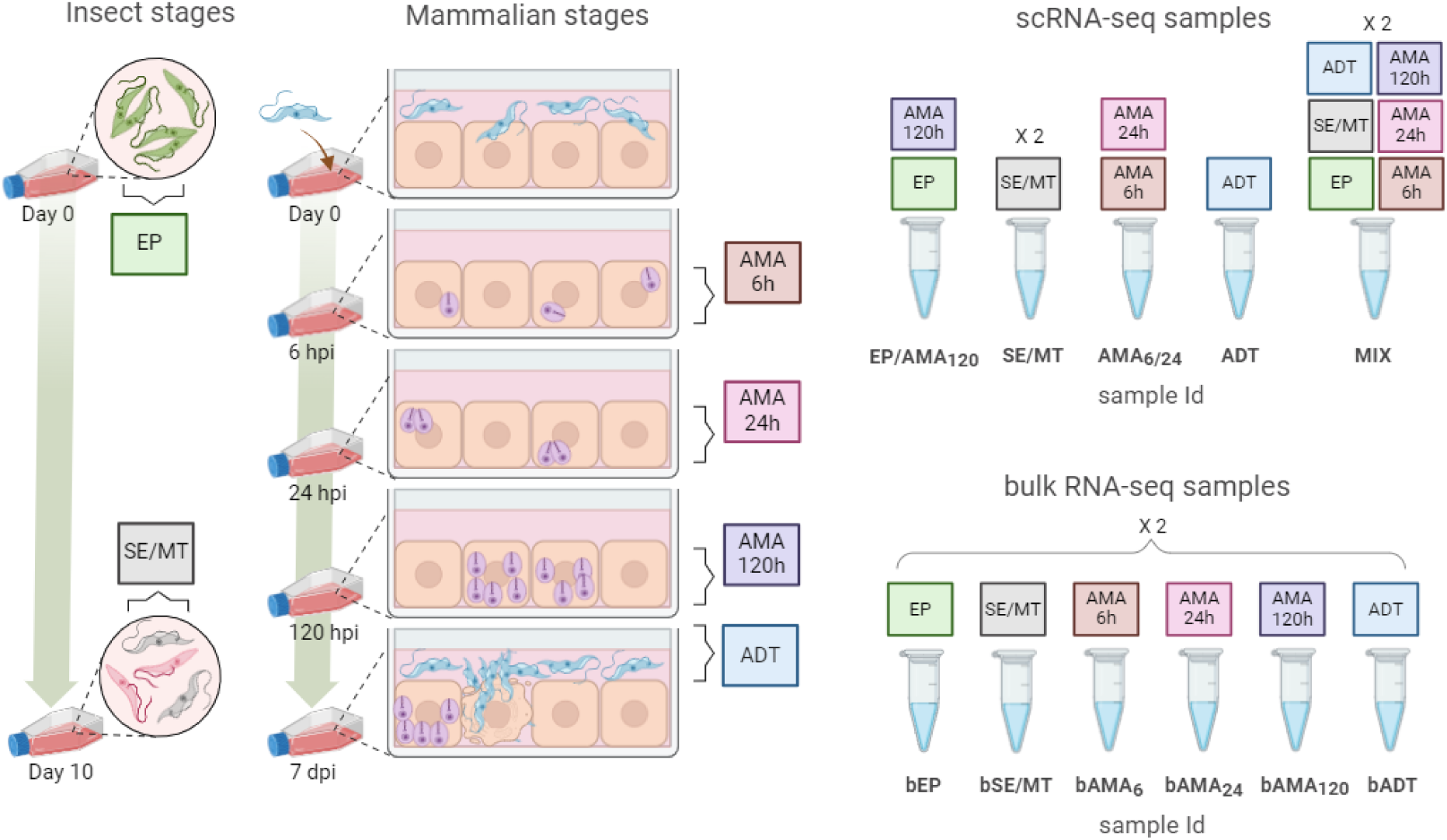
Sampling and library construction. Samples collected from *in vitro* cultures: EP: epimastigotes, collected during the exponential growth phase, SE/MT: a mixture of stationary growth phase epimastigotes and metacyclic trypomastigotes, ADT: amastigote derived trypomastigotes, and AMA: amastigotes isolated from Vero-infected cells at 6, 24, and 120 hours post infection (AMA6h, AMA24h and AMA120h respectively). The sampling times for collecting AMA were based on the findings by Li and colleagues, (2016)^11^. This study revealed significant remodelling of the *T. cruzi* transcriptome within the first 24 hours following invasion, with minimal further changes observed during the 24–72 hpi period. To capture the transition from amastigotes to trypomastigotes, the collecting time of 120 hpi was chosen. The SE/MT sample was collected instead of purifying the metacyclic trypomastigote population, to preserve the natural progression between insect stages, allowing for a more comprehensive analysis of the transcriptome changes that occur during metacyclogenesis. Individual libraries were constructed for SE/MT (two biological replicates) and ADT from single-cell suspensions. Two more libraries were prepared by combining AMA6h with AMA24h (AMA6/24), and EP with AMA120h (EP/AMA120) in equal proportion. A second biological replicate was generated by pooling equal numbers of EP, SE/MT, ADT, AMA6h, AMA24h and AMA120h in a single-cell suspension (MIX). Two libraries (technical replicates) were sequenced for MIX. Additional data for sample preparation can be found in Supplementary Table 2. Figure created in BioRender. Garcia Sanchez, M. (2023) BioRender.com/h07k595.

**Figure 2.**
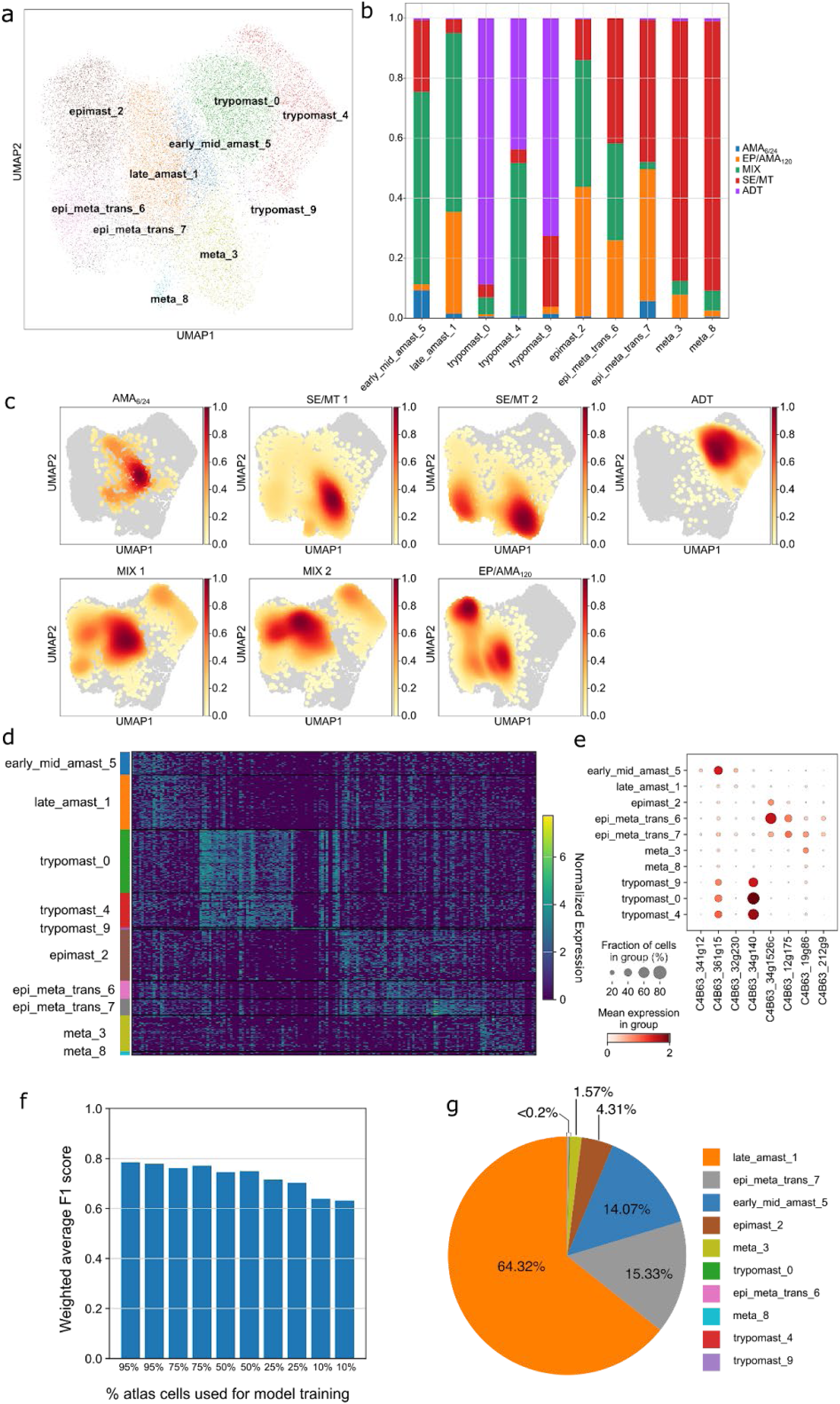
*T. cruzi* atlas overview. **a**, UMAP embeddings of the cells of the *T. cruzi* atlas where every cell is coloured by their cluster identity. **b**, Proportion plot showing the sample origin of the cells across the clusters of the atlas. **c**, UMAP embeddings of *T. cruzi* atlas cells, split by their sample origin, with each cell coloured by the density of cells in that region of the UMAP. Red means more dense regions of cells, white means less dense. **d**, heatmap showing the normalised expression values of the Scanpy derived top 20 marker genes (where available) for each of the clusters in the *T. cruzi* atlas. **e**, dot plot showing the expression of selected marker genes for the clusters in the *T. cruzi* atlas. Dots are coloured by the median expression of the gene in each cluster and the size of the dots shows the percentage of cells (in that cluster) that express the gene. **f**, bar plot showing the weighted average F1 score of scPoli predictions when trained on different percentage subsets of the atlas, against the rest of the cells in the atlas. **g**, pie chart showing the scPoli cluster ID predictions for amastigotes at 48h post infection when trained on the entire *T. cruzi* atlas.

Our first goal, having created the basis for the atlas, was to assess if clusters reflect the different *T. cruzi* life cycle stages that we have captured in our samples. The identified clusters consist of cells with similar transcriptomes and should thus represent cells in the same life cycle stage. We sought to confirm this initially based on our sampling strategy. Fig. 2b shows the sample origin for all the cells in each cluster and Fig. 2c visualises where cells from each sample are located on the UMAP plot. Clusters 0, 4 and 9 contain a high percentage of cells from the ADT sample, indicating these clusters represent amastigote-derived trypomastigotes. Clusters 1 and 2 both mainly originate from the MIX and EP/AMA_120_ samples, indicating these clusters represent either epimastigotes or late-stage amastigotes. Cluster 5 contains many cells from AMA_6/24_ and is adjacent to cluster 1 on the UMAP, but not to cluster 2. Cluster 5 thus likely represents amastigotes in the early stages of development after host cell infection, cluster 1 amastigotes in late infection and cluster 2 epimastigotes. The remaining clusters contain many cells from the SE/MT sample. Clusters 3 and 8 are almost completely made up of SE/MT sample cells thus likely representing metacyclic trypomastigotes. Clusters 6 and 7 contain cells from the EP/AMA_120_ and the SE/MT samples and may represent cells that are transitioning from epimastigotes to metacyclic trypomastigotes, as we generated metacyclic trypomastigotes by prolonged culturing of epimastigotes to trigger differentiation, resulting in a mixed population of epimastigotes and metacyclic trypomastigotes.

### Cluster mapping by marker genes

The above analysis based on our sampling strategy provides a first indication that the parasites are indeed clustered by life cycle stage. However, unsuccessful integration of the datasets could lead to incorrect partitioning of cells into different clusters. Furthermore, interpreting the distances between points on a reduced dimension projection has been shown to be problematic due to distortions when reducing an ambient space with many thousands of dimensions to just two^34^. To increase our confidence in the biological relevance of the clustering we sought to associate marker genes for our clusters with known life cycle markers. The Scanpy-derived top 20 marker genes per cluster (Supplementary File 1), where available, were plotted in a heatmap for all cells in the integrated dataset (Fig. 2d) with specific genes highlighted in Fig. 2e. Substantially increased expression of the trans-sialidases C4B63_34g140 and C4B63_361g15 in clusters 0, 4 and 9 is consistent with these clusters representing the trypomastigote population^35,36^. The predicted amastigote clusters (1 and 5) showed increased expression of the amastigote-associated genes C4B63_32g230, a member of the amastin glycoprotein family^37^ and C4B63_341g12, a lipase associated with early amastigote development^38^. Interestingly, the trans-sialidase C4B63_361g15 also showed increased expression in the early-mid amastigote population, indicating that downregulation of this trans-sialidase occurs later in the trypomastigote to amastigote differentiation compared to C4B63_34g140. TcSMUGL (C4B63_34g1526c) expression, a mucin-type glycoconjugate restricted to the surface of *T. cruzi* epimastigotes^39^, associated clearly with 2, 6, and 7, supporting our assignment of these as epimastigote containing clusters. In line with clusters 6 and 7 representing differentiating epimastigotes, we saw increased expression of C4B63_12g175 and C4B63_212g9, two putative members of the Nodulin-like/Major Facilitator Superfamily, which has been previously associated with the epimastigote to metacyclic trypomastigote transition^18^. Finally, clusters 3 and 8, which we annotated as metacyclic trypomastigotes, show increased expression of C4B63_19g86, encoding CYC2-like cyclin 6, which has been previously shown to be upregulated in metacyclic trypomastigotes^7,16^. Taken together, the marker gene analysis supports our cluster allocation based on our sampling strategy.

### scRNA-seq atlas allows confident life cycle stage mapping

To assess the suitability of our scRNA-seq atlas to serve as a reference to predict cell type labels on an unlabelled dataset, we first tested whether the atlas could be used to correctly annotate a subset of cells from the atlas itself. Using scPoli^40^, a model was trained on subsets of the atlas and used to predict the life cycle stage labels of the rest of the dataset. As model training can be computationally expensive, we trained the scPoli model using increasingly smaller training data sizes to find an optimal trade-off between annotation accuracy and training size. Across all the subsets tested, the weighted average f1-score did not drop below 0.6 (Fig. 2f). Furthermore, the 95% training data subset only returned a slightly higher weighted average f1-score than the 50% training data subset, showing the atlas can accurately label the cells with only half the atlas (Fig. 2f). Variation was observed in the classifying accuracy of the different clusters. The epimast_2 and meta_3 clusters had f1-scores of ∼0.84 when using 95% of the data to train the model on and ∼0.76 when only 5% was used (Supplementary Table 5). Conversely, the epi_meta_trans_7 cluster f1-score with the 95% training subset was ∼0.58 and decreased to ∼0.35 for the 5% training subset (Supplementary Table 5).

As the testing and training datasets are derived from cells that come from similar sequencing runs and were generated in similar ways, this represents a best-case scenario for predicting life cycle stages. As future researchers’ data may be generated through different means or under different conditions, we next aimed to see whether the atlas could be used to correctly annotate cells that were sampled independently from those in the atlas itself. We generated a separate scRNA-seq dataset of amastigotes harvested 48 hours post infection and used a scPoli model trained on the entire *T. cruzi* atlas to predict life cycle stage labels. As these amastigotes lie between the mid and late stages post infection amastigotes, we considered a cluster annotation of late_amast_1 or early_mid_amast_5 as correct and anything else as incorrect. Using this approach, 78.4% of cells were correctly annotated as amastigotes (Fig. 2g), confirming the suitability of our atlas for supervised labelling of *T. cruzi* scRNA-seq data.

### Bulk RNA-seq across all life cycle stages validates the scRNA-seq atlas

To further validate our scRNA-seq atlas, we generated bulk RNA-seq data for key life cycle stages outlined in Fig. 1 (Supplementary Tables 6 and 7). The obtained data showed clear distinction between life cycle samples, and concordance between replicates (Fig. 3a). We next identified marker genes for each sample relative to all other samples (Fig. 3b), as well as for each life cycle stage relative to all other life cycle stages (i.e. all amastigote samples combined) (Fig. 3c). This analysis allowed us to identify highly specific marker genes for each life cycle stage, with 344 genes significantly upregulated in bEP, 510 in bSE/MT, 1,381 in bAMA, and 1,842 in bADT, many of these in line with previously published results (Supplementary File 2). As expected, a substantial proportion of the bADT marker genes encode for multigene family surface proteins, including 413 trans-sialidases from groups I-VIII, 320 mucin-associated surface proteins (MASP), 168 mucins and 44 GP63s. This is in sharp contrast with the identification of only two GP63 and five trans-sialidases from groups I and II as markers for the bSE/MT sample. Well defined markers for the other life cycle stages were also identified such as amastin C4B63_32g230, C4B63_32g229, C4B63_268g12, C4B63_268g18 and C4B63_6g559 for amastigotes and the mucins C4B63_109g84, C4B63_300g114c and C4B63_34g1450c for epimastigotes. Interestingly, we were able to identify specific markers for different stages of amastigote development (Fig. 3b, Supplementary File 3). Chagasin (C4B63_32g1409c) is a marker of the early amastigote population (bAMA_6_) and thus more highly expressed in this population than in any of the others, suggesting that inhibition of cruzipain, or other cysteine proteases, is important at the early stages of amastigote differentiation. Another marker of interest for this population is ATG7 (C4B63_84g61), suggesting a role for autophagy in early amastigotes^41^. The late bAMA_120_, marker genes include one mucin I (C4B63_63g77) and two mucin II encoding genes (C4B63_11g418 and C4B63_63g63), a hexose transporter (C4B63_14g3) and two prostaglandin E synthase-2 genes (C4B63_8g300 and C4B63_50g199).

**Figure 3.**
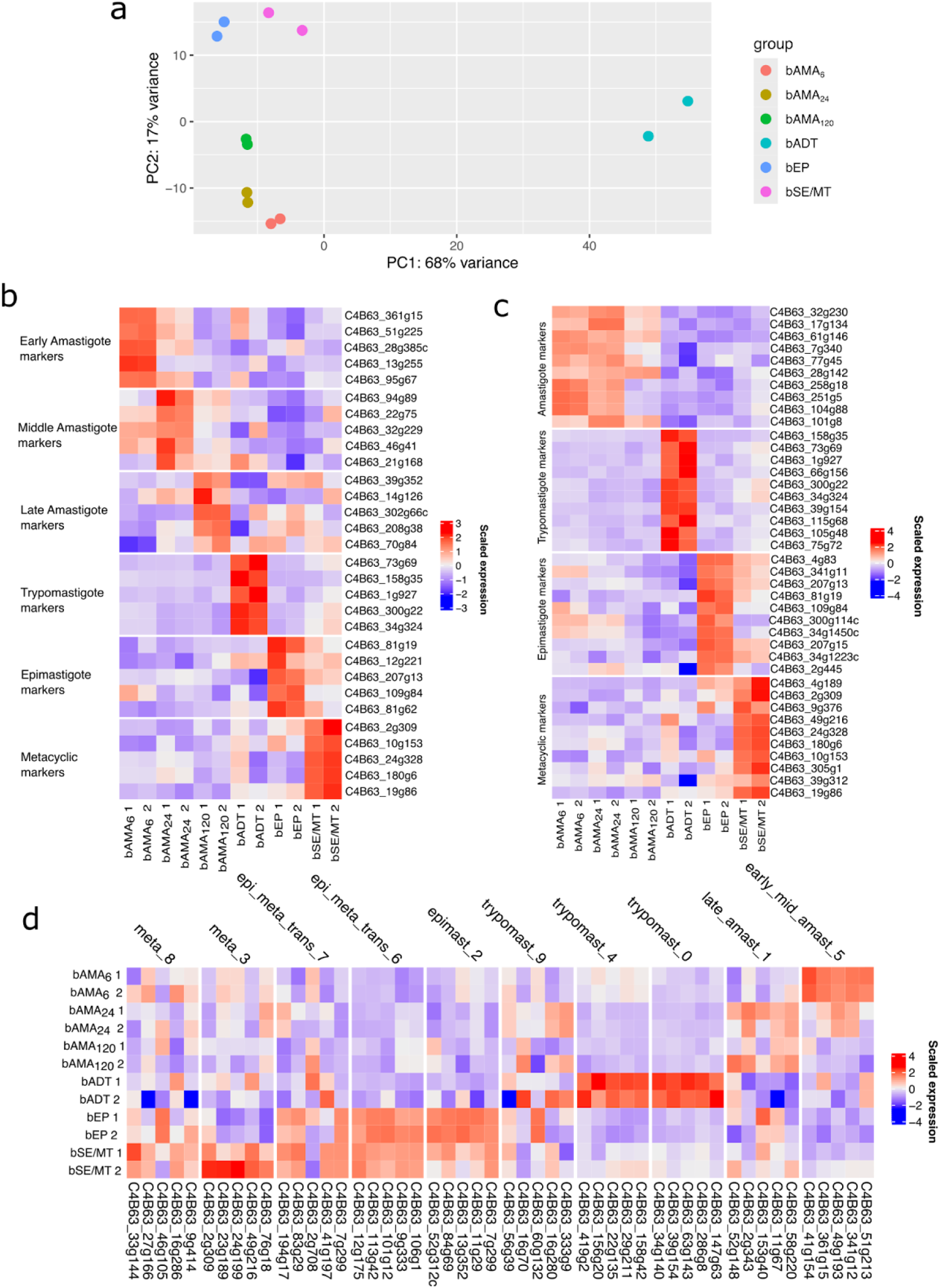
Analysis of bulk RNA-seq datasets of *T. cruzi* life cycle stages. **a**, PCA plot showing the PC embeddings of the bulk RNA-seq datasets, where each dataset is coloured by what *T. cruzi* lifecycle stage is contained within it. **b**, heatmaps showing the scaled, normalized expression of the top 10 marker genes in terms of log2FC for all *T. cruzi* lifecycle stage samples. **c,** the top 10 marker genes in terms of log2FC for the main stages of the *T. cruzi* lifecycle. **d,** heatmap showing the scaled, normalised expression of the top 5 marker genes (in terms of log2FC) for the scRNA-seq datasets clusters across the bulk RNA-seq datasets.

Next, we assessed whether the expression of marker genes derived from our scRNA-seq clusters match up with their respective bulk RNA-seq clusters (Fig. 3d). The markers of the scRNA-seq clusters were generally highest expressed in their equivalent bulk RNA-seq samples across both biological replicates. Some marker genes were highly expressed across multiple life cycle bulk RNA-seq samples, with the markers of single-cell clusters 6 and 7 highly expressed in both the bEP and bSE/MT bulk RNA-seq samples (Fig. 3d), supporting that clusters 6 and 7 represent epimastigote to metacyclic trypomastigote transitionary cells. Markers for cluster 9, which consists of a relatively small number of cells (Fig. 2a) mainly originating from the ADT sample (Fig. 2b), showed less clear alignment with the bulk data, with some genes expressed highly in amastigotes and trypomastigotes, potentially indicating this cluster contains transitional forms between amastigotes and trypomastigotes.

The comprehensive bulk RNA-seq data presented here thus further validates our scRNA-seq cluster assignments and provides a new population level resource to study transcriptomic changes during the *T. cruzi* life cycle.

### UTR mapping

Trypanosomatid mRNA boundaries are defined by the splice leader acceptor site at the 5’UTR start and the poly-adenylation site at the 3’ UTR end. Partial annotation of *T. cruzi* UTRs, in specific life cycle stages, has been carried out^42,43^, but genome-wide annotation is currently not available. Our 3’ enriched scRNA-seq and full-length transcript bulk RNA-seq data from all lifecycle stages allowed us to comprehensively annotate mRNA boundaries (Fig. 4a). The average and median lengths for the 5’UTR are 400bp and 116bp (11,908 transcripts), and 1171bp and 563bp for the 3’ UTR (16,955 transcripts), out of a total of 17,197 transcripts. Our analysis estimates that there are ∼1,500 missing genes in the DM28c annotation^44^ (Fig. 4b, Supplementary Fig. 2a). We also found evidence for ncRNA, with high GC content, but no clear ORF (Supplementary Fig. 2b). Interestingly, we identified ∼18,000 alternative poly-adenylation sites (Fig. 4b). While we saw evidence for lifecycle-stage specific alternative poly-adenylation (Supplementary Fig. 2d), we could not establish this robustly. Finally, we observed two types of reverse strand transcription. In the scRNA-seq data we observed, for some genes, reverse strand expression at the 3’UTR end, in the opposite direction of the main transcript (Fig. 4b). In the bulk RNA-seq data we detected reverse strand transcripts covering the full length of the primary transcripts (Supplementary Fig. 2c). Enrichment analysis of the top 500 genes with the highest level of antisense transcripts revealed enrichment of genes involved in cell adhesion (p < 0.001), in particular MASPs (Bonferroni 2.35e-4).

**Figure 4.**
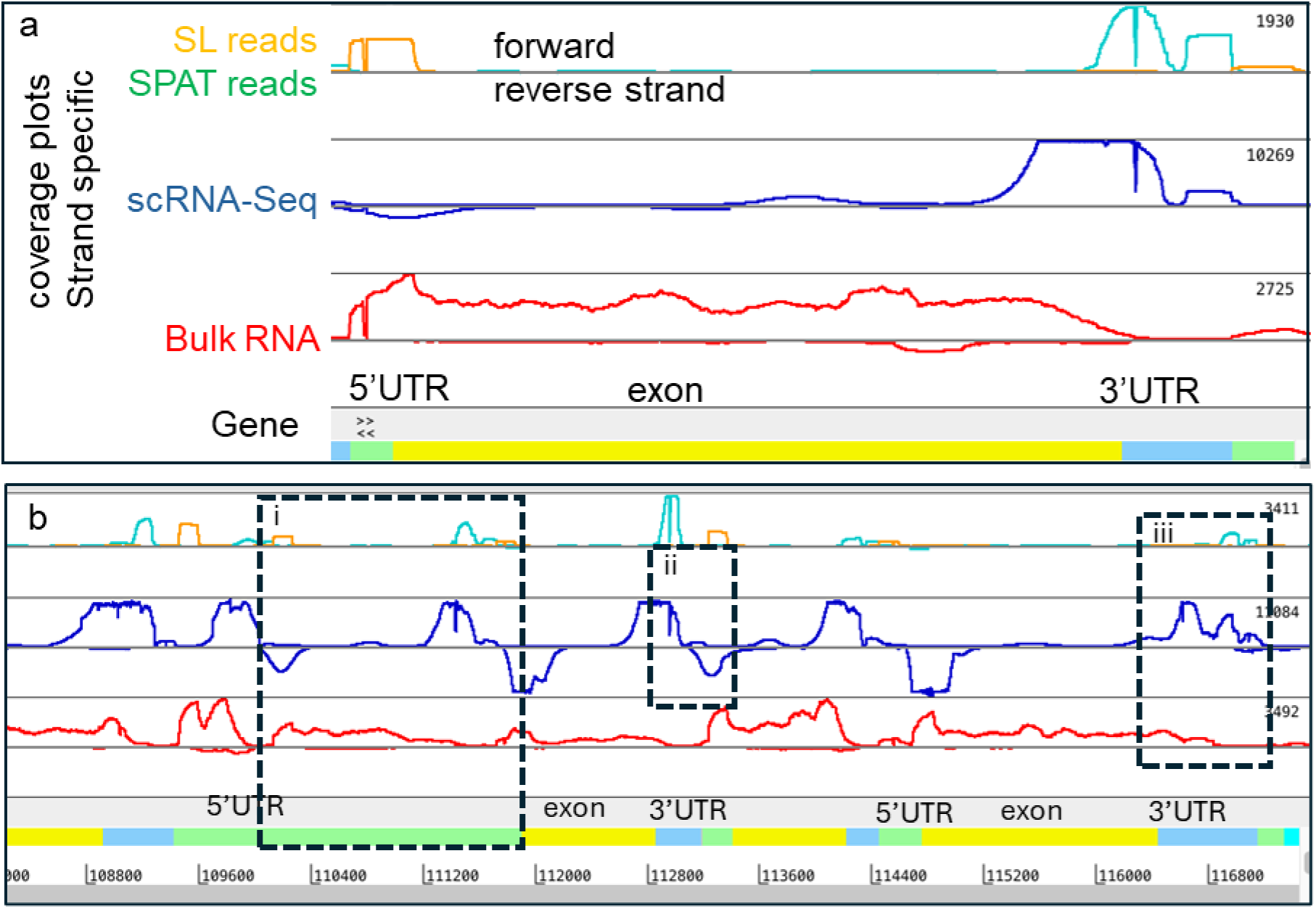
Annotation and analysis of UTRs. **a**, example of gene (C4B63_1g67-t42_1) annotated with UTRs (green 5’, blue 3’). Read coverage is shown for scRNA-seq (blue) and bulk RNA-seq (red). Splice leader (SL) is identified in bulk RNA-seq data (orange), poly-adenylation sites are identified in the scRNA-seq data using SPAT (green). Numbers on right indicate read numbers at top of scale. **b**, Region of chromosome 1, with three genes (annotation as in a). (i) evidence for unannotated genes, with SL and SPAT in 5’ UTR of downstream gene, (ii) scRNA-seq evidence for reverse strand transcript at 3’UTR, (iii) example of alternative poly-adenylation sites (two blue peaks in scRNA-seq, and confirmed with lower coverage in bulk RNA-Seq).

### Differentiation trajectory analysis identifies genes involved in metacyclogenesis

scRNA-seq provides a key advantage over bulk RNA-seq when studying cells undergoing transitions between different life cycle stages, as bulk RNA-seq does not have the resolution to identify the signal coming from these cells, which may be very few in a given dataset. We can see this lack of resolution when looking at the expression of our epimastigote to metacyclic trypomastigote transitionary clusters markers on our bulk RNA-seq data (Fig. 3d). We sought to identify genes whose expression changes across transitions and thus may play a key role in differentiation. As we have clusters containing epimastigotes, metacyclic trypomastigotes and those that appear to be transitioning between the two, we utilised trajectory inference (TI) to identify genes that are associated with the epimastigote to metacyclic trypomastigote transition (Fig. 5a-c). This analysis identified 231 genes whose expression peaked early in the transition, 571 genes that peaked during the middle stages and 121 genes that peaked at the end of the differentiation process (Fig. 5d, Supplementary File 5), providing a rich dataset to study metacyclogenesis. As examples (Fig. 5e), we found early expression of genes involved in amino-acid catabolism, such as L-threonine dehydrogenase (LDH) (C4B63_42g129), 2-amino-3-ketobutyrate coenzyme A ligase (KBL) (C4B63_32g217) and proline racemase (C4B63_52g138). Genes that peak during the transition (“middle peak”) are particularly relevant with respect to the differentiation process. Within this category, we identified C4B63_34g1526c, which encodes a mucin TcSMUG-L surface protein known to be restricted to the surface of epimastigotes, where it is thought to play a role in attachment to the insect midgut epithelium^39,45^. C4B63_12g175, a member of the Nodulin-like/Major Facilitator Superfamily that we discussed above as a marker for clusters 6 and 7, also peaked during the transition, in line with a role during differentiation. Several DNA repair proteins, namely RAD51, RAD50 and RAD54 (C4B63_6g507, C4B63_6g508 & C4B63_28g105), were also found to peak in expression during the middle phase of metacyclogenesis. As expected, known markers for metacyclic trypomastigotes peaked towards the end of the transition, including the surface protease GP63 (C4B63_16g183)^9,46^, the cruzipain inhibitor chagasin (C4B63_32g1409c)^47^ and citrate synthase (C4B63_4g184)^48^.

**Figure 5.**
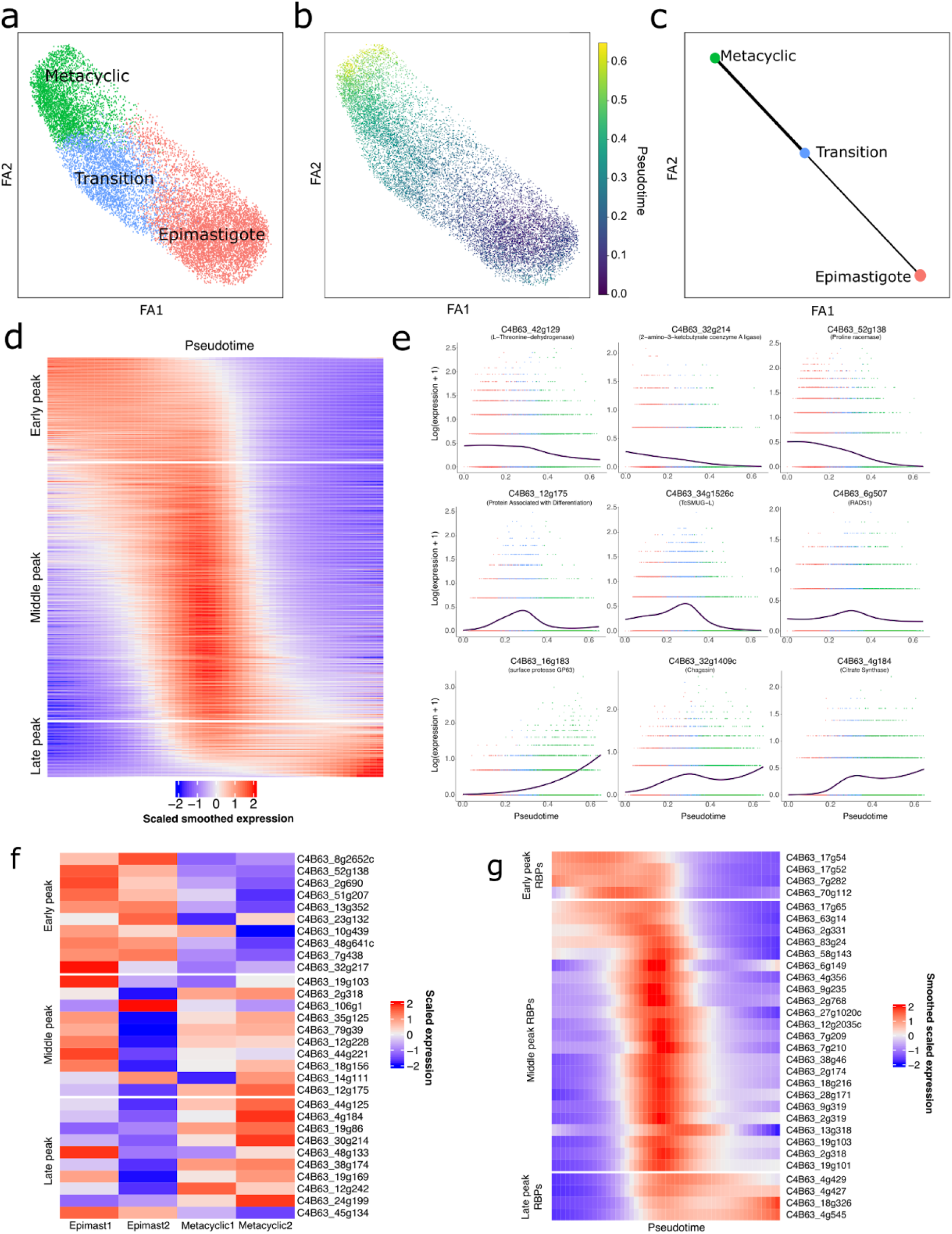
Trajectory analysis of epimastigote to metacyclic trypomastigote transition. **a**, Force atlas 2 (FA2) embeddings of the epimastigote, metacyclic trypomastigote, and transitionary cells coloured by cluster identity and **b**, PAGA-derived pseudotime. **c**, PAGA graph on the FA embeddings, with the dots representing the epimastigote, metacyclic trypomastigote and transitionary clusters and the thickness of the lines showing how connected the clusters are, as determined by PAGA. **d**, heatmap showing the scaled smoothed expression values (across 50 sampled points across pseudotime) for the genes whose expression significantly changed over the metacyclogenesis axis. Genes are ordered along the Y axis by their peak expression across the metacyclogenesis process. **e**, plots showing the log1p transformed expression of selected genes whose expression significantly changed over metacyclogenesis pseudotime. Cells are coloured by their cluster identity and the lines represent the smoothed expression of the gene across pseudotime. **f**, heatmap showing the scaled, normalised expression of the top 10 genes (in terms of Log2FC) for the early, mid and late peaking genes, whose expression significantly changed over metacyclogenesis pseudotime, on the epimastigote and metacyclic trypomastigote bulk RNA-seq samples. **g**, heatmap showing the scaled smoothed expression values (across 50 sampled points across pseudotime) for the RNA binding protein-encoding genes whose expression significantly changed over the metacyclogenesis axis. Genes are ordered along the Y axis by their peak expression across the metacyclogenesis process.

To assess the validity of the markers as being important in the epimastigote to metacyclic trypomastigote transition, we looked at the expression of the markers in the bulk RNA-seq data (Fig. 5f). Interestingly, while the genes that defined the early and late phases of differentiation were highest expressed in the epimastigote and metacyclic trypomastgote bulk RNA-seq datasets respectively, no clear pattern was seen for the genes identified as peaking in the middle of the process, supporting the power of scRNA-seq over bulk approaches to identify transitionary clusters and extract their gene signatures.

Regulation of mRNA stability and half-life play a key role in kinetoplastid gene expression control. RNA-binding proteins (RBPs) are trans-acting elements that engage with cis-acting regulatory sequences in the untranslated regions of mRNAs, mediating mRNA processing, stabilisation, subcellular localization, and translation efficiency^49^. Increasing evidence suggests that the expression levels of specific RBPs are closely associated with the regulation of subsets of mRNAs, influencing parasite differentiation in both *T. cruzi* and *T. brucei* ^11,50^. We therefore examined our pseudotime dataset specifically for RBPs (Fig. 5g). This revealed RBPs previously associated with differentiation. For example, C4B63_19g103 peaked during the middle phase of the epimastigote to metacyclic trypomastigote transition. Its ortholog in *T. cruzi* CL Brener (TcCLB.507093.220, TcUBP1) has been shown to play a role in differentiation of *T. cruzi* epimastigotes into metacyclic trypomastigotes under nutritional starvation conditions^50–52^. C4B63_4g545, encoding RNA-binding protein 4, showed increased and sustained expression over pseudotime; peaking in the late phase of the transition, in alignment with the increased expression of its CL Brener orthologue (TcCLB.508901.20) in metacyclic trypomastigotes^7^. The expression of C4B63_18g326, encoding the zinc-finger protein ZFP1, also peaked late in the transition process. A role for ZFP1 in *T. brucei* bloodstream form differentiation has been suggested^53^, and upregulation of this RNA-binding protein (RBP) in the stationary stage of *T. cruzi* epimastigote growth curve has been previously described^54^. We thus identified RBPs with known association to differentiation. In addition, we identified a large number of less well-characterised RBPs that associate with pseudotime during metacyclogenesis and may play important roles in gene expression control during differentiation. In total, we identified four early peak RBPs, 23 middle peak RBPs and four late peaking RBPs. These genes are of key interest for further study of the regulation of metacyclogenesis.

### Distinct trypomastigote populations

Our single-cell analysis revealed heterogeneity within life cycle stages. Within the trypomastigote population, we identified two substantial sub-populations (clusters 0 and 4), in addition to the small cluster 9. We isolated clusters 0 and 4 to identify markers that distinguish these trypomastigote sub-populations (Supplementary File 4). GO term analysis of the markers for cluster 0 shows high expression of genes required for protein synthesis and key metabolic pathways (Fig. 6a), whereas the key markers for cluster 4 are mainly trans-sialidases and GP63 surface proteins (Fig. 6b, c, d). Our data thus reveals the presence of two transcriptionally distinct trypomastigote populations.

**Figure 6.**
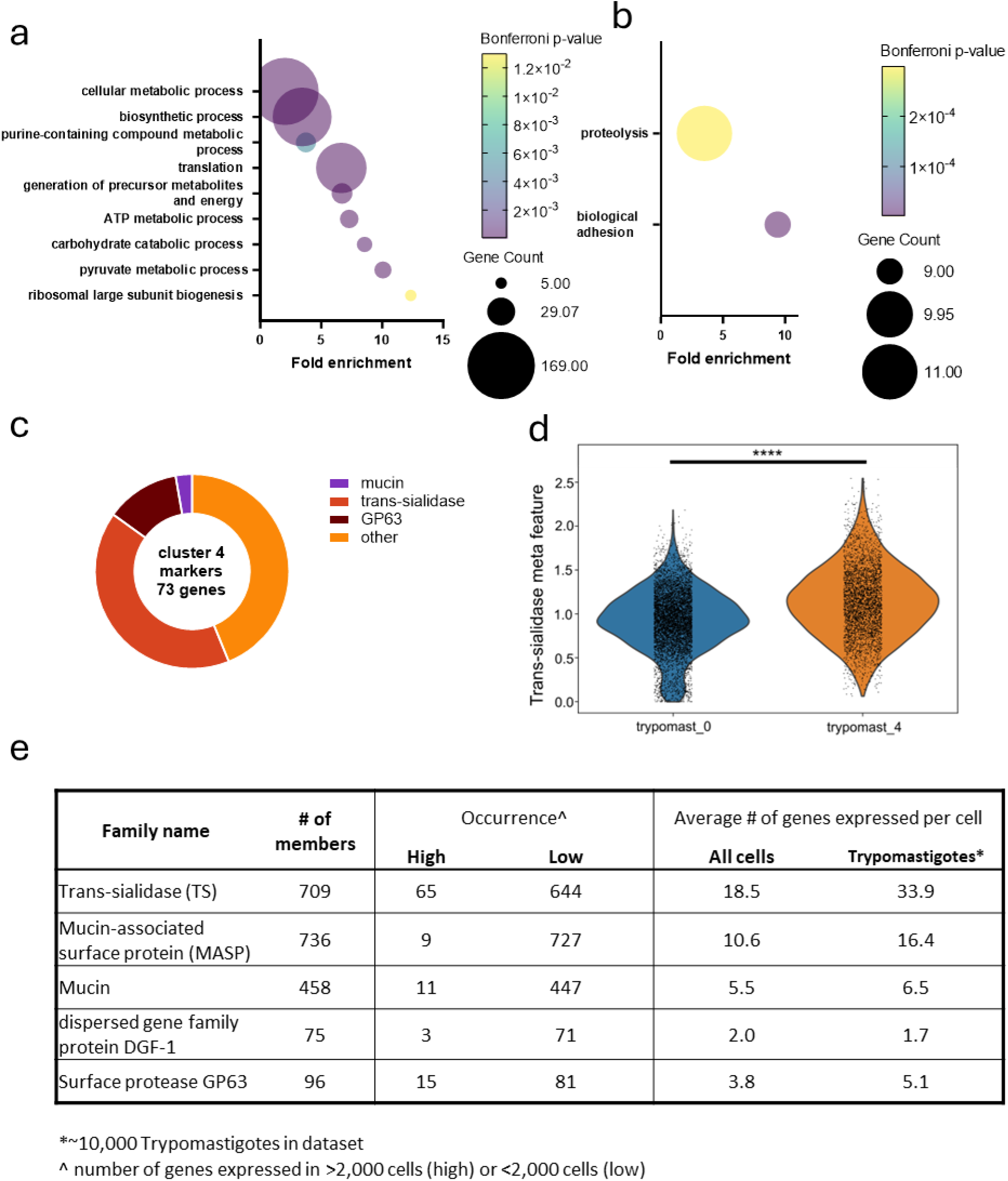
Two distinct trypomastigote populations. **a**, GO molecular function terms enrichment analysis for marker genes for cluster 0. Fold enrichment, number of genes found for each term and adjusted p-value (Bonferroni) are displayed. GO terms containing substantial overlap in genes are not displayed. **b**, GO molecular function terms enrichment analysis for marker genes for cluster 4. **c**, pie chart for protein family distribution of the 73 cluster 4 marker genes. **d**, violin plots showing expression levels of all detected trans-sialidases combined for each cell. The trypomast_4 cluster shows significantly higher trans-sialidase expression levels compared to trypomast_0 (two-sided independent t-test). **e**, multi-gene family repertoires from single cell data. Relevant histograms are shown in Supplementary figures 3, 5 and 5.

### Cell specific multi-gene family repertoires

The *T. cruzi* genome comprises multiple large gene families such as trans-sialidases (TS), GP63s and mucins. At a population level, many members of these families are co-expressed, depending on the life cycle stage. We assessed the extent of co-expression at the single cell level in 10,359 trypomastigotes (ADT), which express a large number of multi-gene family surface proteins. For the TS we found that a subset of 65 genes (out of 709) was expressed in >2000 cells, whereas the remainder were expressed in many less cells (Fig. 6e, Supplementary Fig. 3a). On average, 33.9 TS were expressed in each trypomastigote (Fig. 6e, Supplementary Fig. 3b). In individual cells, the number of commonly expressed TS relative to number of rare TS was normally distributed, with most cells expressing an equal number of common and rare TS (Supplementary Fig. 3c). These data indicate that each trypomastigote expresses a different repertoire of TS, made up of a small set of high-occurrence TS and a diverse set of rarer ones. These observations applied across the three ADT clusters (Supplementary Fig. 3d). Linking occurrence to sequence similarity revealed a small cluster of closely related Group IV trans-sialidases with high occurrence (Supplementary Fig. 3e). Indeed, 99% of trypomastigotes express at least one of these closely conserved Group IV TS. Other TS groups had a much higher frequency of low occurrence genes. Similar analyses on the other surface protein families showed that individual mucin and MASP genes typically have a low occurrence profile, whereas the distribution for GP63s is more akin to the TS family (Fig. 6e, Supplementary Figs. 4 & 5). Interestingly, a subset of closely related GP63 genes, representing TcGP63 Group 9^55^, is expressed in most cells, whereas other GP63 sub-groups showed more sporadic expression.

## Discussion

Single-cell transcriptomic approaches provide an unprecedented view of the heterogeneity in cell populations. Here we present the first scRNA-seq data for *Trypanosoma cruzi*, across a comprehensive set of life cycle stages, including intracellular amastigotes, resulting in a single-cell transcriptomic atlas for this important human pathogen. We extensively validated the assignment of cell types in the atlas, using known marker genes and bulk RNA-seq. The atlas shows distinct clusters for all life cycle stages, with associated marker genes, as well as clusters representing transitionary cells.

To validate life cycle assignments in the single-cell atlas, we compared all life cycle stages against each other using bulk RNA-seq for the first time. In addition to confirming correct life cycle stage assignment in the single-cell atlas, this bulk RNA-seq dataset provides a new resource, identifying highly life cycle stage specific markers, including markers specific for the different amastigote timepoints.

Combining our scRNA-seq and bulk RNA-seq data allowed us to accurately map the 3’ and 5’ UTRs of many genes, providing improved UTR annotation of the *T. cruzi* genome. Interestingly, we found many alternative poly-adenylation sites. Alternative poly-adenylation (APA) is common in eukaryotes, and has been described previously in *Trypanosoma brucei*^56–58^. Our data shows that APA is common in *T. cruzi*, and as in other organisms it likely plays a key role in controlling mRNA stability and translation efficiency^59^. Further work is required to understand the extent of life-cycle specific APA.

Antisense transcripts have been shown to play a role in gene expression regulation in trypanosomes^60–62^. Our data reveals a previously underappreciated complex landscape of antisense and reverse strand transcripts, potentially revealing an important new layer in gene regulation.

An important use for cell atlases is to serve as a reference to map cells in new datasets. Here we show that our atlas can be used reliably for life cycle stage mapping, with a median f1-score of 0.756 across the 10 clusters (Fig. 2f, g). To put this into perspective, the weighted average f1 score when applied to mammalian datasets, which have more accurate transcriptomic references, ranges between ∼0.5 and ∼1.0 depending on the dataset^40^, supporting the quality and utility of our atlas for life cycle stage mapping.

Our single-cell approach led to identification of two subpopulations in the amastigote-derived trypomastigote population (Fig. 4), with one population (cluster 0) apparently more metabolically active, as indicated by the over-representation of marker genes associated with metabolism, and the other population (cluster 4) characterised by expression of classical trypomastigote surface genes. These two subpopulations are in accordance with previously observed morphologically and functionally distinct trypomastigote subpopulations^2,63,64^, and the marker genes we identified provide a stepping stone to further investigate the underlying molecular determinants of these distinct trypomastigote populations.

For organisms with complex life cycles such as *T. cruzi*, a key power of the single approach lies in resolving individual cells progressing through the multiple life cycle transitions, thereby allowing temporal resolution of gene expression and identification of genes whose expression is associated with differentiation. Not all life cycle transitions are fully covered by our atlas; for example we did not obtain data for the metacyclic trypomastigote to amastigote transition. We did obtain samples to study the tissue culture trypomastigote to amastigote to trypomastigote transitions, but trajectory analysis could not unambiguously identify transitory cells, although we did identify markers for the amastigotes at different timepoints post infection. Based on comparison with bulk RNA-seq data, cluster 9 may contain transitionary forms, but we could not confirm this. Potentially our sampling approach did not capture a substantial number of cells undergoing differentiation, or differentiation is rapid, meaning that only very few cells are at the same stage at any one point in time, rendering identification difficult.

Trajectory analysis for the epimastigote to metacyclic trypomastigote (metacyclogenesis) was successful. We identified a substantial number of genes whose expression was associated with pseudotime (Fig. 5). This analysis was validated by comparison with bulk RNA-seq and known markers and revealed a large number of novel genes associated with epimastigote to metacyclic trypomastigote differentiation. We focused our attention on RNA-binding proteins, which play a key role in gene expression regulation in trypanosomatids^49,65^. In total we identified 31 RBPs whose expression was associated with pseudotime, with of particular interest the 23 RBPs whose expression peaked during the transition. These RBPs are prime candidates to be regulators of gene expression during metacyclogenesis and thus for further study.

Population level studies have demonstrated life-cycle dependent expression patterns for multi-gene families in *T. cruzi*. Here we explore for the first time the heterogeneity of expression patterns at the single cell level, and our finding that the repertoire of key surface gene-families varies between individual cells expands our understanding of the parasite’s approaches to dampen the immune response. In addition to the smokescreen generated by the large number of surface antigens, and their release^66,67^, the cell-specific surface antigen repertoires mean that most antibodies can likely only neutralise a subset of the parasites, further delaying an effective immune response. Importantly, some sub-groups of closely-related genes, such as the group IV trans sialidases and Group 9 GP63 genes, show a very high occurrence at the single cell level, pointing to a core function and making them potential candidates for vaccine approaches. While various antigens of group IV TS TC13 (C4B63_98g20) have been unsuccessfully explored to generate a productive vaccine^68–70^, our data indicates it may be worth exploring other antigens in this family, in particular those that are highly conserved between all members in Group IV.

This first single-cell transcriptomics dataset for *T. cruzi* provides an important resource for the research community and opens new research avenues to study *T. cruzi* life cycle heterogeneity and differentiation. Furthermore, it will guide future scRNA-seq experiments, to look in more depth at specific differentiation steps and explore alternative biological contexts, such as the impact of host cell type on the parasites and parasites isolated *ex vivo*.

## Funding

This research was funded by the Medical Research Council through New Investigator award MR/T030283/1 to MDR and their Precision Medicine DTP (MRC grant number: MR/N013166/1) and in part, by Wellcome Awards 104111/Z/14/ZR (to TDO) and 203134/A/16/Z (to MDR).

## Materials and methods

### 2.1. Parasite culture

*T. cruzi* parasites from the Silvio strain (MHOM/BR/78/Silvio; clone X10/7) subclone A1 were used in this study^71^. Epimastigotes were grown at 28 °C in RTH/FBS [RPMI 1640 medium (Sigma, St Louis, MO, USA) supplemented with 0.4% (w/v) trypticase peptone (BD, Le Pont de Cleix, France), 25 μM hemin (Sigma, St Louis, MO, USA), 17 mM Hepes (Formedium LTD, Hunstaton, UK) pH7.4, and 10% (v/v) heat-inactivated foetal bovine serum (FBS, Gibco, Paisley, UK)]. The cultures were maintained in exponential growth by continuous passages of 3×10^5^ cells/ml every three days.

Metacyclogenesis was induced by nutritional starvation, as previously described^20^. Briefly, epimastigotes were transferred to a culture flask containing fresh RTH/FBS to a density of 3×10^5^ cells/ml and were maintained at 28 °C without media change. The relative percentage of metacyclic trypomastigotes to stationary phase epimastigotes was determined by microscopic examination of parasites. On day 10 of culture, the highest percentage of parasites that displayed a metacyclic trypomastigote-like shape (∼50%) was observed, aligning with findings reported for the Medellin strain (TcI)^10^.

Intracellular amastigotes were grown in Vero cells, as described by MacLean et al.^20^. In brief, metacyclic trypomastigote-rich cultures were incubated with a monolayer of Vero cells (ECCAC 84113001) at a multiplicity of infection (MOI) of 10 overnight at 37 °C 5% CO_2_ in Minimum Essential Medium (MEM, GlutamaxTM Supplement) (Gibco, Paisley, UK) supplemented with 10% FBS. After overnight infection, extracellular parasites were removed, and the Vero cell monolayer was washed three times with serum-free MEM, followed by addition of fresh MEM 10% FBS. Infected flasks were maintained at 37°C 5% CO_2_ and the medium was replaced at 72 hours post-infection (hpi). At seven days post-infection (dpi) amastigote-derived trypomastigotes emerging from infected Vero cells were harvested and used to set up new infections at MOI 1.5.

### 2.2. Sample collection

Epimastigotes (EP) were collected in the exponential growth phase, from 2 days-old cultures initiated with 3×10^5^ cells/ml. A mixture of stationary phase epimastigotes and metacyclic trypomastigotes (SE/MT) was obtained from a 10 days-old culture initiated with 3×10^5^ epimastigotes/ml. Amastigote-derived trypomastigotes (ADT) were harvested from the supernatant of infected Vero cells at 7 dpi. EP, SE/MT and ADT were put in falcon tubes and left undisturbed for 1 h (at 28 °C for EP and SE/MT, and 37 °C 5% CO2 for ADT). To avoid collecting dead parasites and co-purified Vero cells, the supernatant, containing viable and motile cells was collected. Parasites were pelleted and washed once in ice-cold phosphate-buffered saline (PBS) by centrifugation at 1,350 x g for 5 min at 4 °C.

Amastigotes (AMA) were obtained from Vero cells infected with ADT and washed at 6 hpi as indicated above. At the time points selected (6, 24, and 120 hpi), intracellular AMA were isolated from the host cells following the protocol described by Dumoulin et al. (2020)^72^, with minimum modifications. MOI 1.5 was used for 120 hpi samples and MOI 10 for 6 and 24 hpi samples. Infected monolayers were washed with ice-cold PBS and scraped into 3 ml PBS. Infected cells suspensions were lysed using the gentleMACS^TM^ dissociator with gentleMACS^TM^ M tubes (Miltenyi Biotec, Bergisch Gladbach, Germany) and the “Protein” protocol. The ensuing lysates were passed through a PD-10 desalting column (Cytiva, Little Chalfont, UK) equilibrated with PBS. AMA were eluted in 3.5 ml ice-cold PBS and were washed three times in 15 ml PBS by centrifugation at 1,350 x g for 5 min at 4 °C, to remove host cell RNA and debris.

The resulting pellets containing EP, SE/MT, ADT, and the three time points for AMA (AMA_6h_, AMA_24h_ and AMA_120h_) were resuspended in ice-cold PBS and kept on ice. For bulk RNA-seq experiments, 2×10^7^ parasites per sample were pelleted and frozen on dry ice until RNA extraction. For scRNA-seq experiments, parasites were diluted in PBS to 1×10^6^ cells/ml.

### 2.3. Cell viability assessment

Aliquots from all parasite stages diluted at a density of 1×10^6^ cells/ml were stained for 5 min at room temperature with SYTOX™ AADvanced™ Dead Cell Stain (Life Technologies, Eugene, OR, USA) following manufacturers’ recommendations. A total of 20,000 events per sample were acquired using a CytoFLEX flow cytometer (Beckman Coulter, Indianapolis, IN, USA) set to excitation at 488 nm and emission at 647 nm wavelengths. The percentage of viable cells was determined with the CytExpert v2.4 software (Beckman Coulter). Parasites killed with three cycles of freezing and thawing were used as positive control for SYTOX AADvanced staining.

### 2.4. Bulk RNA-seq – RNA extraction, library preparation and sequencing

Total RNA was extracted using RNeasy Mini Kit (Qiagen, Hilder, Germany) following the manufacturer’s protocol, with an on-column DNase digestion step with the RNAse-Free DNase Set (Qiagen). Two biological replicates for each life cycle stage were prepared and sent to Novogene Co., Ltd (Cambridge, UK) for rRNA removal, library construction and sequencing. RNA quantity and integrity were measured with an Agilent 2100 bioanalyzer using the RNA Nano 6000 Assay Kit (Agilent Technologies, Santa Clara, CA, USA) and by agarose gel electrophoresis. A total of 1 µg total RNA per sample was used as input material for the library preparation. Ribosomal RNA was depleted from total RNA and strand-specific libraries were generated using NEBNext® UltraTM RNA Library Prep Kit for Illumina® (New England BioLabs, Ipswich, MA, USA) following manufacturer’s recommendations. Index codes were added to attribute sequences to each sample.

Quality of the library was assessed by Agilent 2100 bioanalyzer followed by quantification by qRT-PCR. The different libraries were pooled according to the effective concentration and the target amount of data – 6 Gb raw data per sample – and were sequenced by the Illumina NovaSeq 6000 (Illumina Inc., San Diego, CA, USA). Paired-end 150bp reads were generated.

### 2.5. Single-cell RNA-seq – library preparation and sequencing

Single-cell RNA-seq experiments were performed using Chromium Single Cell Gene Expression solution (10X Genomics, Pleasanton, CA) according to the manufacturer’s guidelines. Single-cell suspensions of 16,500 cells in PBS were prepared for SE/MT and ADT. AMA_6h_ was combined with AMA_24h_, and EP was combined with AMA_120h_ in equal proportion, in two cell suspensions of 16,500 cells each. Two technical replicates were collected for the SE/MT sample. A second biological replicate was prepared in a separate experiment by pooling equal quantities of EP, SE/MT, ADT, AMA_6h_, AMA_24h_ and AMA_120h_ for 21,000 cells in total. For this pool (labelled MIX) two technical replicates were collected (Fig. 1). Single-cell libraries were generated using the Chromium Next GEM Single Cell 3ʹ Reagent Kit v3.1 and the Chromium Next GEM Single Cell 3ʹ Reagent Kit v3.1 - Dual Index (10X Genomics) for the first and second biological replicates respectively. Briefly, single-cell suspensions were loaded onto a Chromium Controller instrument in a Chromium Next GEM Chip G (10X Genomics) to obtain single-cell Gel beads in EMulsion (GEMs). Barcoded cDNA was obtained from GEMs and amplified by PCR in a PCRmax Alpha Cycler 1 thermal cycler (Cole-Parmer, Saint Neots, UK). End repair and A-tailing, adapter ligation, post-ligation cleanup with SPRIselect (Beckman Coulter, Brea, CA, USA), and sample index PCR and cleanup steps were performed as per the manufacturer’s instructions. Final library sample index PCR cycle parameters were selected based on quantification of barcoded cDNA samples with a Qubit 2.0 (Life technologies, Carlsbad, CA, USA) using the Qubit dsDNA HS Assay Kit (Life technologies, Eugene, OR, USA): SE/MT samples were amplified by PCR with 16 cycles while the rest of samples were amplified with 14 cycles. Single-cell libraries were sent to Novogene Co., Ltd for sequencing. Libraries were quantified by Qubit and quality was tested in an Agilent 2100 Bioanalyzer before sequencing on the Illumina NovaSeq 6000 platform using the PE150 strategy for 20 Gb raw data per sample.

### 2.6. Bioinformatics analysis

#### 2.6.1. Creating reference transcriptomes for mapping

The *Trypanosoma cruzi* Dm28c 2018 genome and annotation files were edited to remove rRNA, tRNA, ncRNA and pseudogene sequences. The 3’ UTRs annotations of the *Trypanosoma cruzi* Dm28c 2018 ^73^ were extended by 2500 base pairs as described in Briggs et al., (2021)^30^. Two combined reference genomes were created with 10X Genomics Cell Ranger v6.0.0. Combined reference genome 1 was made up of: 2500 base pair 3’ UTR extended *Trypanosoma cruzi* Dm28c 2018, *Trypanosoma cruzi* maxicircle kDNA sequence and the grch38 human reference and combined reference genome 2 was made up of: 2500 base pair 3’ UTR extended *Trypanosoma cruzi* Dm28c 2018 and the *Trypanosoma cruzi* maxicircle kDNA sequence.

#### 2.6.2. Mapping and counting of life cycle bulk RNA-seq samples

All samples were mapped against combined reference genome 1 using STAR on default parameters ^74^. Count matrices for the samples were generated using featureCounts ^75^, where multi-mapping and multi-overlapping genes were not counted, the minimum overlapping bases was one and paired reads were treated as a single fragment, not two individual reads.

#### 2.6.3. Processing and analysis of bulk RNA-seq samples

The genes whose expression was less than 10 in any 11 of our 12 samples were removed from the dataset as well as the kDNA maxicircle genes. To identify marker genes of the different life cycle stages/samples, each life cycle stage/sample was contrasted against all other life cycle stages/samples using the results() command in DESeq2 ^76^, with the list values for the stage of interest being one and the list value for the other stages being one divided by the number of stages in that side of the contrast. Genes were identified as markers of the life cycle stage if they were significantly differentially expressed (i.e. they had a Benjamini and Hochberg adjusted p-value < 0.05), were upregulated (Log2 fold change > 0.5) in a given life cycle stage/sample, compared with the other samples/stages.

#### 2.6.4. Mapping and counting scRNA-seq reads

For each sample, reads uniquely mapping to annotated genes on the created references were counted and assigned to a cell using the Cell Ranger Count function (Cell Ranger v6.0.0 - Default settings). Samples SE/MT and ADT were mapped against combined reference genome 2, while AMA_6/24_, EP/AMA_120_ and MIX samples were mapped against the combined reference genome 1. Around 30% of the reads mapped to the reference in Cell Ranger, which uses the STAR read mapper. To understand what the missing 70% were, we repeated the mapping with a less stringed mapper (BWA)^77^. This increased the mapping ∼ 80%. We observed that 60% of the mapped reads located to five contigs that all contain rRNA genes. As shown in Supplementary Figure 6, STAR (blue line) does not map as many reads to rRNA region as BWA (black line). We did not investigate the mapping differences further, however, to understand why so many reads of rRNA are captured, we looked at SPAT for possible evidence of polyadenylation of rRNA and homopolymer tracks of As. As show in Supplementary Figure 6, the number of SPAT reads is less than 0.5%, compared to the coverage height. However, we find several homopolymers of at least 6bp, which are probably sufficient to capture rRNA during the 10X Chromium reaction.

#### 2.6.5. Quality control, processing and integration of scRNA-seq samples

Count data output from Cell Ranger was loaded into an AnnData object for analysis using Scanpy ^78^. Cell quality was assessed by looking at the total number of genes captured per cell, the percentage of reads mapping to the kDNA maxicircle and the percentage of reads mapping to the human genome (for the samples that were mapped against combined reference 1, containing the human genome). Cells with high numbers of genes detected were removed as they may be doublets, cells with low number of genes detected were removed as they may be dying cells or droplets which only contain ambient RNA and cells with high percentage mapping to the kDNA maxicircle were removed as they may represent dying cells^79^. Cells with a high percentage of reads mapping to the human genome were removed as we wanted to only capture individual parasite cells for analysis. The threshold values used for cell selection for each dataset are given in Supplementary Table 4.

The count matrix of the AnnData ^80^ objects for each sample were normalised through the following process. A cell’s counts were divided by the total sum of counts for the cell, multiplied by 1000, 1 was added to each value of the count matrix (to remove 0 values), and then finally, the counts were transformed by natural logarithm. As some of the samples contain homogenous populations of cells (e.g. the trypomastigote sample, which just contains trypomastigotes), no filtering out of genes was performed in order to retain genes which may uniquely define these subsets of these seemingly homogenous populations.

The individual AnnData objects were then concatenated to make a single AnnData object and the normalised count matrix of all the genes and cells was stored in the raw layer of the AnnData object. The top 2500 most variable genes (in terms of normalised dispersion) were found for the concatenated dataset; these genes will be used for the scaling, dimensionality reduction, integration and clustering analyses outlined below. The genes associated with the kDNA maxicircle were removed from the top 2500 most variable genes list to ensure the levels of kDNA gene expression were not driving the clustering or dimensionality reduction.

The effect of the total number of genes per cell was regressed out of the count matrix in the active layer of the AnnData object before the data was scaled to a mean of 0 and unit variance. Any scaled values above 10 were clipped to 10. PCA was performed on the scaled and regressed matrix, with the first 50 principal components (PCs) being calculated. The data was integrated using BBKNN ^81^, with batch key being the metadata column containing which sample each cell came from and default values being used apart from the number of PCs which was chosen to be 17, based on the variance ratio plot of the first 50 PCs. The BBKNN neighbour graph was used as the basis for creating a Uniform Manifold Approximation and Projection (UMAP) ^82^ embeddings of the data and assigning cells to clusters through the Leiden algorithm ^83^ at a resolution of 0.7.

#### 2.6.7. Analysis of epimastigote to metacyclic trypomastigote transitions in scRNA-seq data

The Epimastigote_2, Metacyclic_3 and Epi_meta_trans 7 and 8 clusters were subsetted out into their own object. All subsequent analysis was performed on this subsetted datasets. The data was rescaled and reclustered in Scanpy before pseudotime analysis with PAGA ^84^ was carried out. Cells with PAGA pseudotime values more than 0.65 were removed from the dataset. The data was then converted to SingleCellExperiment ^85^ format for fitting of the general additive models using tradeSeq ^86^, across the PAGA pseudotime. For both datasets, the models were fitted over 7 knots. An association test was then carried out to identify what genes have significant expression changes over the pseudotime axis, where significant genes were those with a log2 fold change more than 0.25, a Bonferroni adjusted p-value of less than 0.05 and expression in at least 10% of cells in the dataset.

To generate the smoothed expression plot sorted by expression peak, we took the genes identified by tradeSeq as being DE and got their smoothed expression from the predictSmooth function. The smoothed expression values were then scaled before they were Fourier transformed using the FitWave function from the metR library. The genes were then ordered by peak across pseudotime.

#### 2.6.8 Reference mapping using the *T. cruzi* atlas

The following training to test data proportion sets were created: 0.95:0.05, 0.75:0.25, 0.5:0.5, 0.25:0.75 & 0.05:0.95. The training sets were then used to train a scPoli model and predict the cell type of the test dataset. This process was repeated once more before weighted average f1-scores were calculated. ScPoli was run using mainly default parameters apart from an embedding dimension of 7 and the condition keys being the sequencing run and sample IDs when training on the atlas training set.

#### 2.6.9 Annotating an unrelated scRNA-seq run using the *T. cruzi* atlas

A new scRNA-seq dataset was generated from amastigotes isolated from 48 hpi cultures. Sample collection and library preparation followed the procedures described in sections 2.2 and 2.5, except that the cultures of Vero infected cells were washed at 16 hpi rather than 6 hpi. A scPoli model was then trained on the entire *T. cruzi* atlas and used to predict the cell type label of the 48 hpi amastigotes.

#### 2.6.10 Gene family analysis

The gene ID of the members of the five gene families were obtained from the current annotation. Raw UMI counts were exported from the scRNA object in Python and associated to the genes. Counts of one UMI were excluded. Analysis and visualisation was performed in R.

To generate the Gephi plots, we first compared the amino acid of all family members against each other with Blastp (parameters: -max_hsps 1000 -num_alignments 1000 -word_size 2 - soft_masking false). The output was taken, and a Perl script transformed the data into the Gephi format^87^. The script also generated a global identity score, by normalising the alignment length to the length of the sequences. In Gephi, the Fruchtermann Reingold method was used to group the genes (nodes) and the Edge Weight (representing the glocal identity) to trim the edges.

#### 2.6.11 UTR annotation

To annotate the UTR we first remapped all the data with bwa mem (default parameters). To define the splice leader reads (SL), from all the bulk RNA-Seq data, we selected read that contained at least 15 bases of the splice leader sequence (AGTTCAGACGTGTGCTCTTCCGATCTGTTTCTGTACTATATT), had a mapping score of at least 5 and that part should be softclipped in the alignment (so not map). We defined a 5’UTR, if there were at least five of those reads supporting the call, it was less than 2.5kb apart from the start codon of the exon and we preferred the “AG” to cover the start of the 5’UTR. We largest the largest 5’UTR. As input, we used the Dm28c2018 annotation from TyriTrypDB, version 68^44^.

For the 3’UTR we used peaks2UTR^88^, which uses *s*oft-clipped polyA-tail *t*runcation (SPAT) reads, which cover the poly-adenylation site, from all the 10X scRNA-Seq data. As input the used the annotation with the 5’UTR, parameters: --no-strand-overlap --do-pseudo -f --max-distance 8000. We are currently in contact to upload that annotation into TriTrypDB.

#### 2.6.12 UTR analysis

For manual inspection and detection of the UTR with used BAMview from Artemis^89^ With the transcriptomics evidence we found new transcripts, confirmed findings and generated the figures. To count the number of alternative 3’UTR end sites, we used SPAT reads with at least 7bp going into the polyA tail and have 41 reads evidence.

To find evidence of expression on opposite strand, we used featureCounts with the parameters: -Q 5 -g ID -t exon -T 8 -p -s 2, basically counting just reads over exons, minimum mapping score and mapping strand specifically. To obtain the top 500 genes with evidence of “reverse strand expression” we generate the ratio with of read mapping on the strand of the exon or the inverse. The top 500 list was used for enrichment analysis on TriTryDB.org.

##### Software versions

STAR version 2.7.11a

FeatureCounts version 2.0.6; Peaks2UTR 1.4.0; BWA 0.7.17-r1188

Analyses in Python were carried out on Python version 3.8.16. Specific package versions for Python analysis are as follows: anndata – 0.9.1, h5py – 3.8.0, leidenalg – 0.10.1, matplotlib – 3.7.1, numpy – 1.24.3, pandas – 2.0.3, pip – 23.0.1, scanpy – 1.9.3, scvi-tools – 0.20.3, scArches – 0.5.10, seaborn – 0.12.2, scipy – 1.19.1, scikit-learn – 1.2.2, umap-learn – 0.5.3, bbknn – 1.6.0, fa2 – 0.3.5, igraph – 0.10.6, pytorch-lightning - 1.9.5

Analyses in R were carried out on R version 4.1.0. Specific package versions for R analysis are as follows: singlecellexperiment – 1.14.1, tradeSeq – 1.6.0, metR – 0.15.0, ComplexHeatmap – 2.8.0

## Notes

### Competing Interest Statement

The authors have declared no competing interest.

### Summary of Updates

Additional information provided regarding UTR annotation (Figure 4). Additional analysis included with respect to cell-surface protein repertoires (Figure 6 and supplementary).

