## Supplementary Information for "The *Trypanosoma cruzi* cell atlas; a single-cell resource for understanding parasite population heterogeneity and differentiation"

**Supplementary Fig 1**. Flow cytometry profiles showing the percentage of viable cells for all the samples collected

A

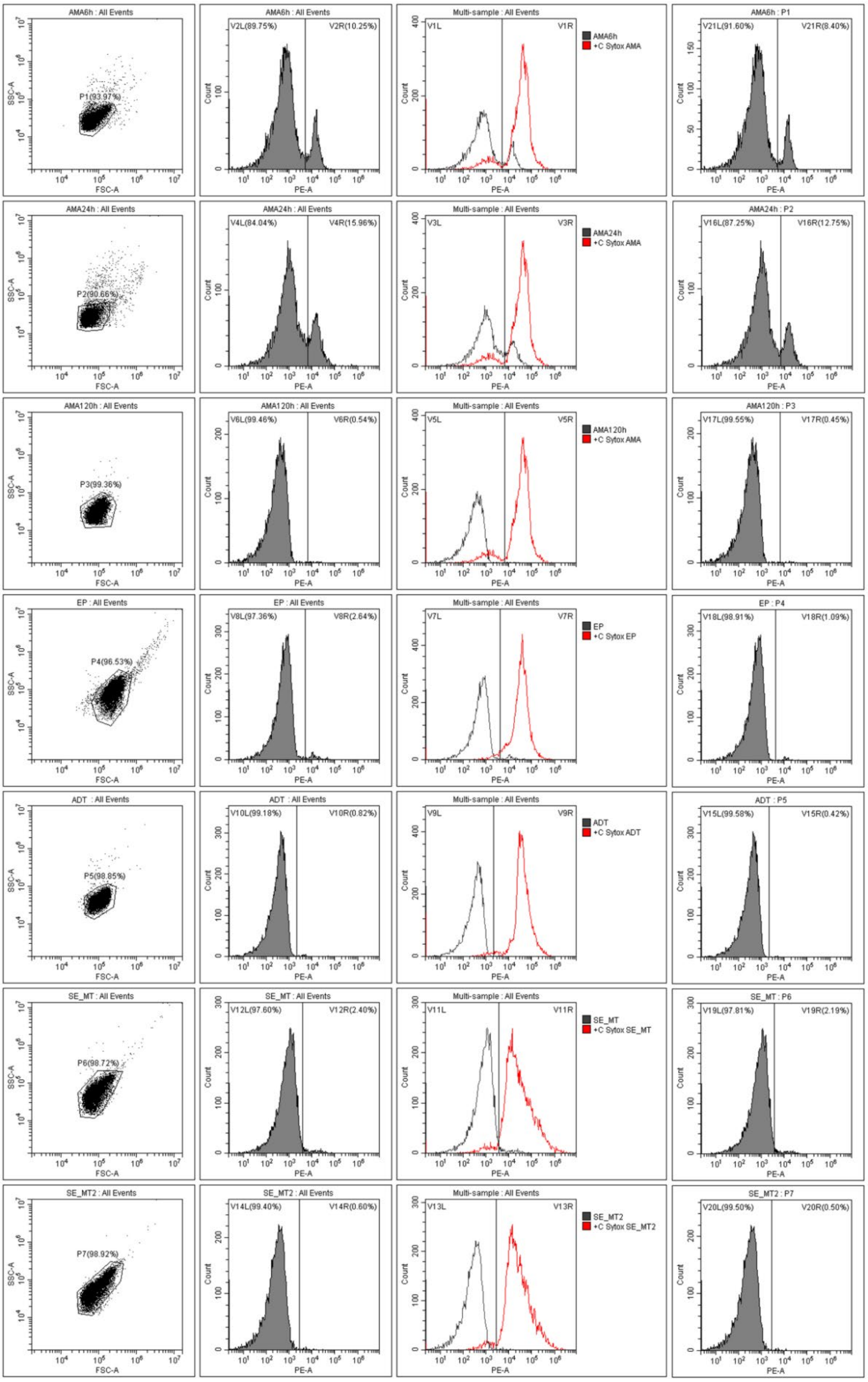

B

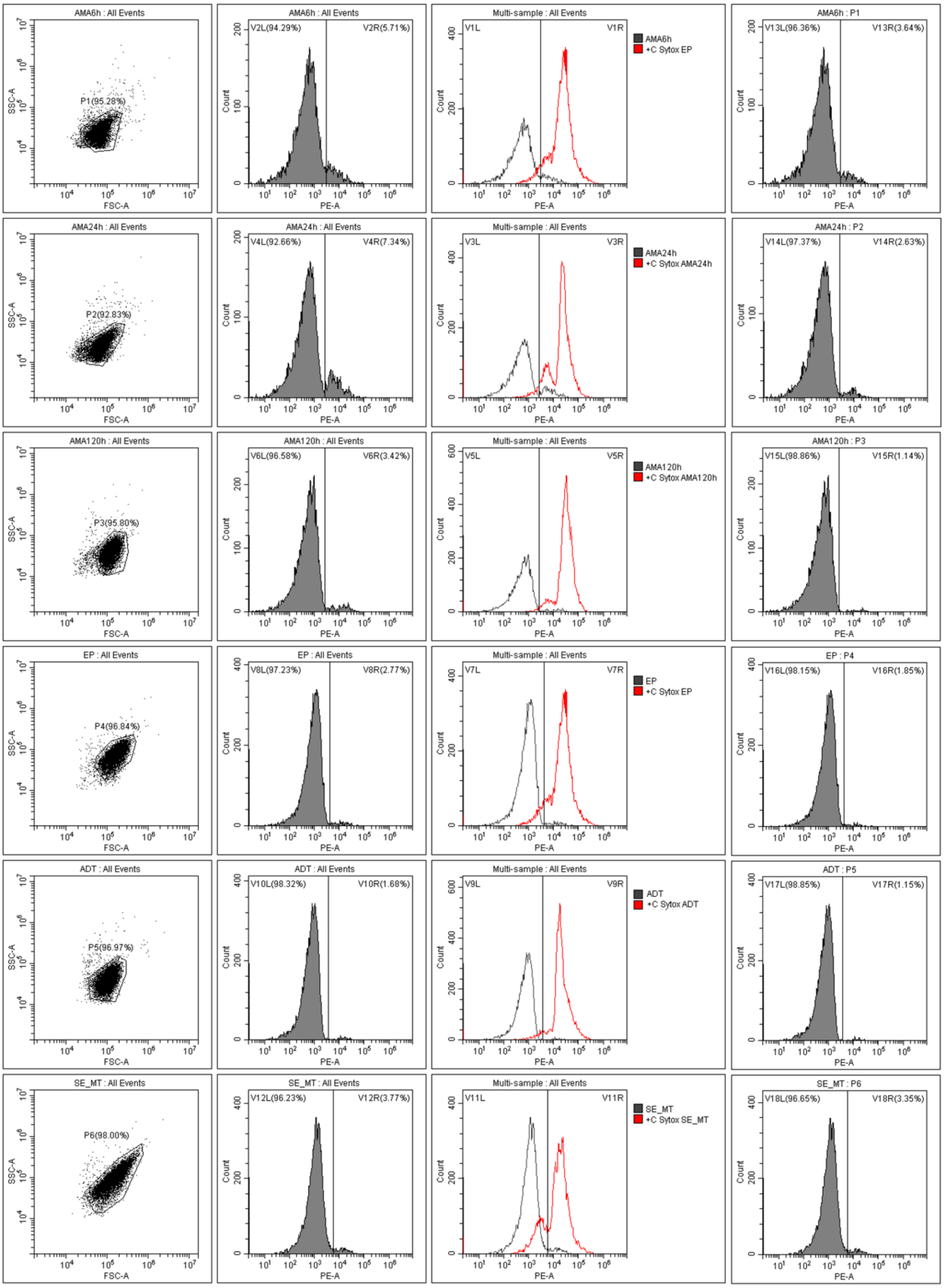


The profiles shown in panel **(A)** correspond to the parasites used for the scRNA-seq samples AMA6/24, EP/AMA120 and SE/MT and the first biological replicate of bulk RNA-seq samples. Those in panel **(B)** were used for the scRNA-seq samples MIX and the second biological replicate of bulk RNA-seq samples.

AMA: amastigotes collected at 6, 24, and 120 hpi (AMA6h, AMA12h and AMA120h respectively); EP: epimastigotes; ADT: amastigote derived trypomastigotes; SE/MT: mixture of stationary phase epimastigotes and metacyclic trypomastigotes obtained by nutritional starvation of EP; C+ Sytox (red line): parasites from the different life cycle stages killed with three cycles of freezing and thawing, used as positive control for SYTOX AADvanced staining. The fourth column shows the profiles of the cells gated in the first column to discard cell debris and clumps.

**Supplementary Fig 2**. Visualisation of transcriptomics data in Artemis

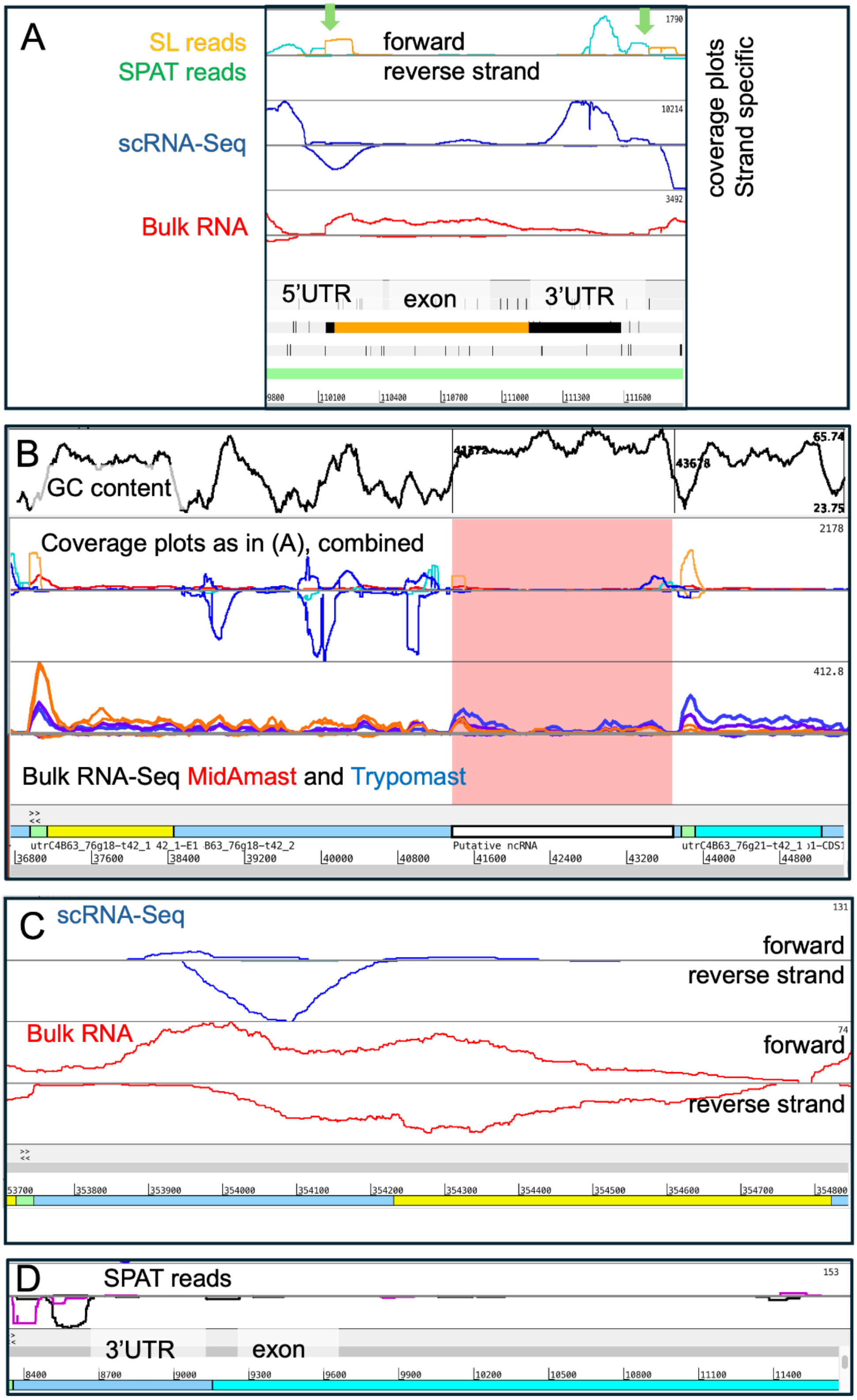


(A) In this long 5' UTR of C4B63_1g48, there is evidence of SL and SPAT (green arrows). The resulting open reading frame (ORF) of nearly 1000 bp has a perfect protein match (100% identity) against BCY84_06787 [*Trypanosoma cruzi* Dm28c 2017]. The red bulk RNA-Seq coverage plot shows evidence of expression, as well as the blue scRNA-Seq coverage plot. (B) Potential new ncRNA. On chromosome 76, in the 3'UTR of C4B63_76g18, we find evidence for a novel transcript. In the bulk RNA-Seq we see evidence for different expression levels between MidAmast (bAMA35, red) and Trypomast (bADT, blue). It was not possible to build a gene model, and although this could be a pseudo gene, due to the high GC content and the lack of clear ORF, we speculate that this is a ncRNA. (C) Example of reverse strand expression of mucin (C4B63_11g381) genes. The gene is on the reverse strand, however on the forward strand, there is clear evidence of "reverse" transcription. Those could be ncRNA that are a further layer of transcriptional control, yet to be explored. Of the top 500 genes with evidence of transcription of the reverse strand (just using the exon, not UTR), 120 are of unknow function, 74 are MASPs, 26 DGF-1 and 24 mucins TcMUCII. (D) Example of alternative 3'UTR end in metacyclic trypomastigotes (violet) and amastigotes (black) in the gene C4B63_1g1402, a retrotransposon hot spot (RHS) protein. The UTR of is ~170bp longer in the metacyclic trypomastigote sample.

All coverage plots were filtered for a mapping score of at least 5 to exclude repetitive mapping reads.

**Supplementary Figure 3**. Single cell expression patterns of the 709 trans-sialidase genes.

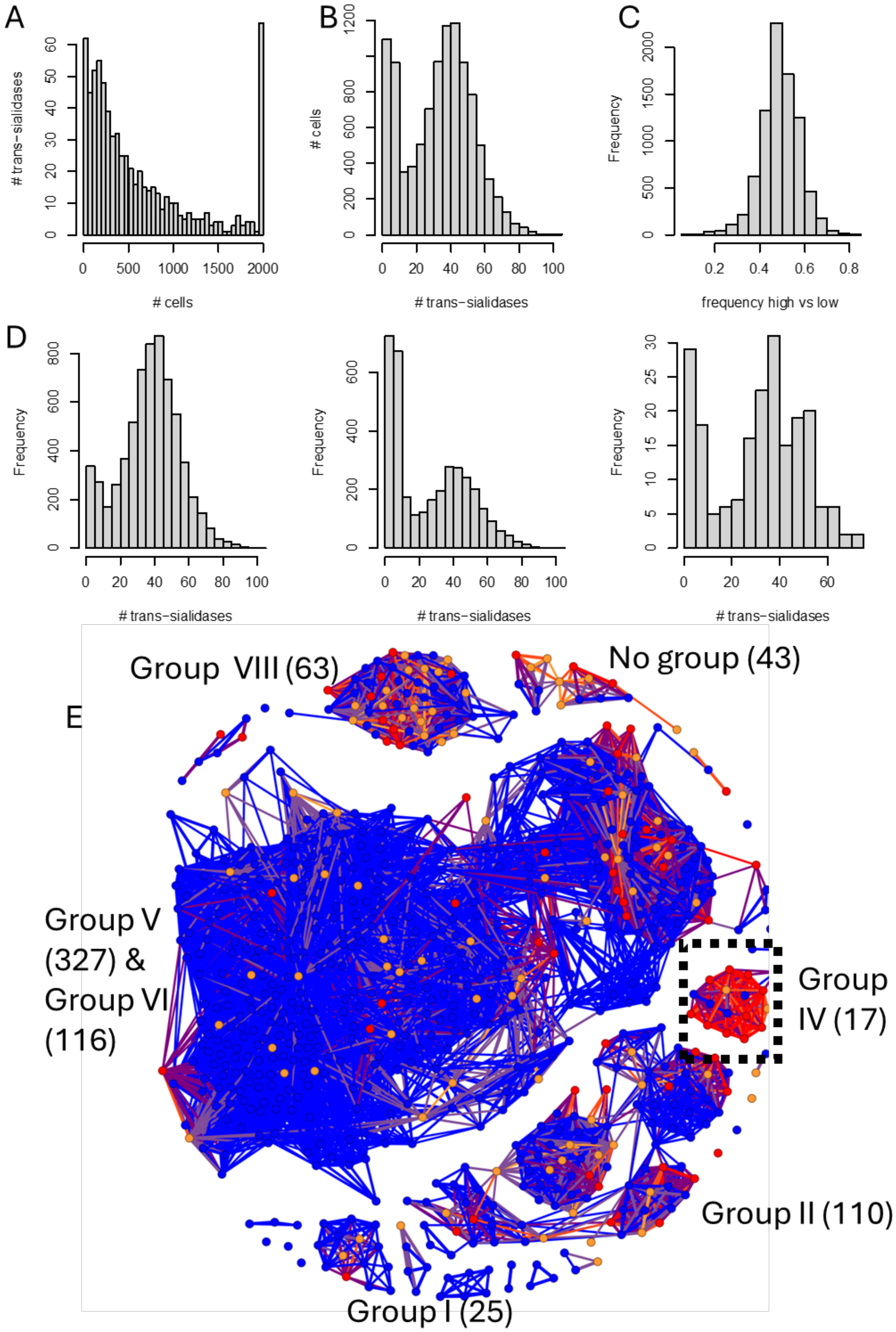

(A) Histogram of number of TS genes (Y-axis) expressed against number of cells expressing these (x-axis). Cut off at 2000 (65 TS). (B) Number of TS expressed per trypomastigote. (C) Frequency of high-expressed genes relative to total (of TS expressed in >20 cells). (D) same as B, but split for the three trypomastigote clusters; clusters 0, 4 and 9, from left to right. (E) Gephi network. Each dot represents a TS, colored by occurrence, in (A), red >2000, orange 1000-2000, blue < 1000 cells. TS are connected by global sequence similarity score > 61%. As indicated in the text, Group IV TS (black dotted box) TS occur in > 99% trypomastigotes (with at least 20 TS expressed). Five members that have > 85% identity to each other are C4B63_34g140, C4B63_109g60, C4B63_109g65, C4B63_34g137, C4B63_98g20. TS groups with more than 15 members are labelled (labels missing for Group III – 7 members & Group VII 2).

**Supplementary figure 4.** Single cell expression patterns of the 736 mucin-associated surface proteins (MASP)

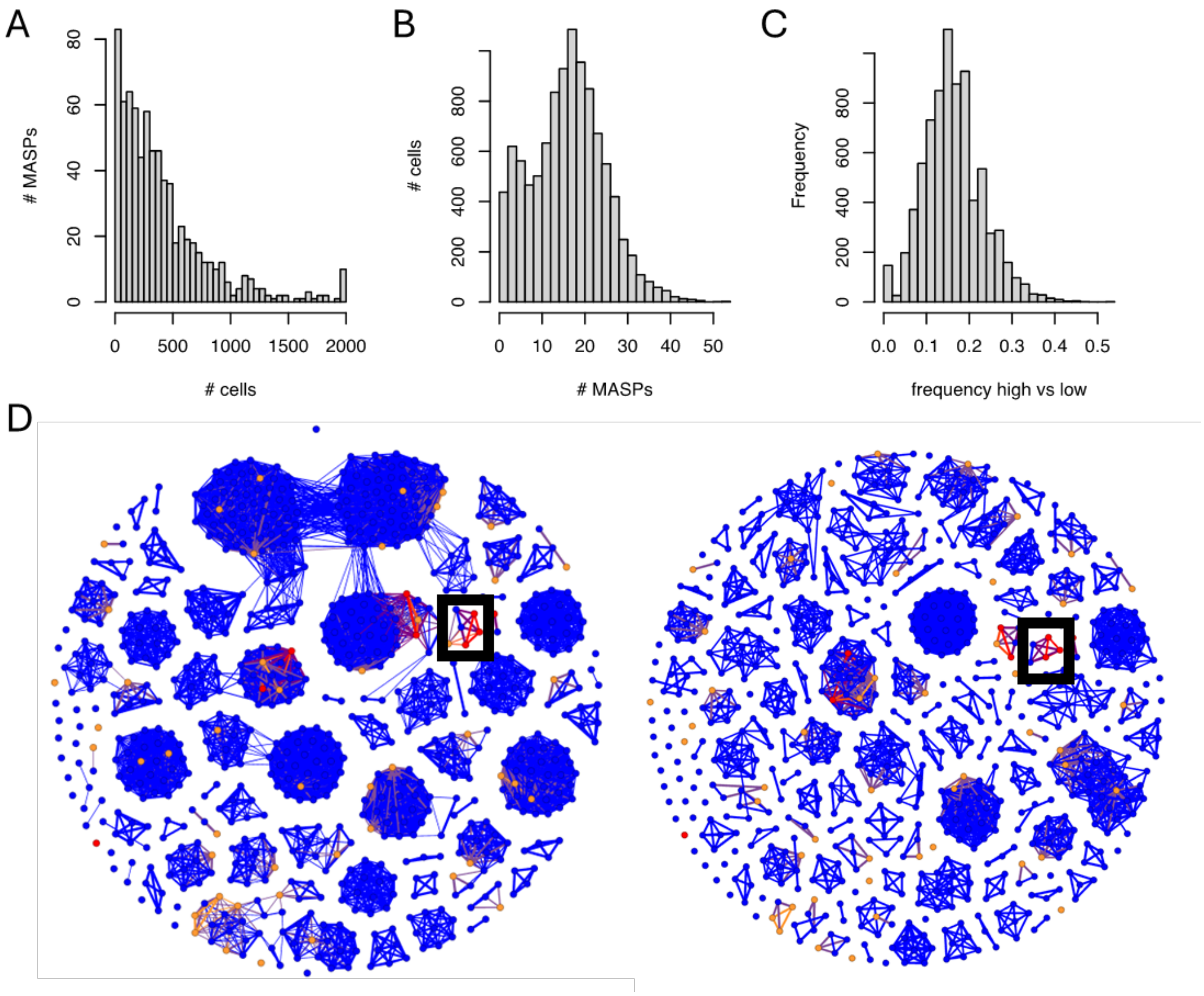


(A) Histogram of number of MASP genes (Y-axis) expressed against number of cells expressing these (x-axis). Cut off at 2000 (9 MASPs). (B) Number of MASPs expressed per trypomastigote. (C) Frequency of high-expressed genes relative to total (of MASPs expressed in >20 cells). (D) Gephi as in Supplementary Figure 3. Left identity cuf-off: >23%, right: >61%. Overall, the MASP gene families seem to be more diverse than the TS. Black box highlights three genes (C4B63_147g63, C4B63_196g38 and C4B63_24g26) and occur in 9773, 8865 and 4984 cells, respectively.

**Supplementary figure 5.** Single cell expression patterns of other multi-gene family surface genes.

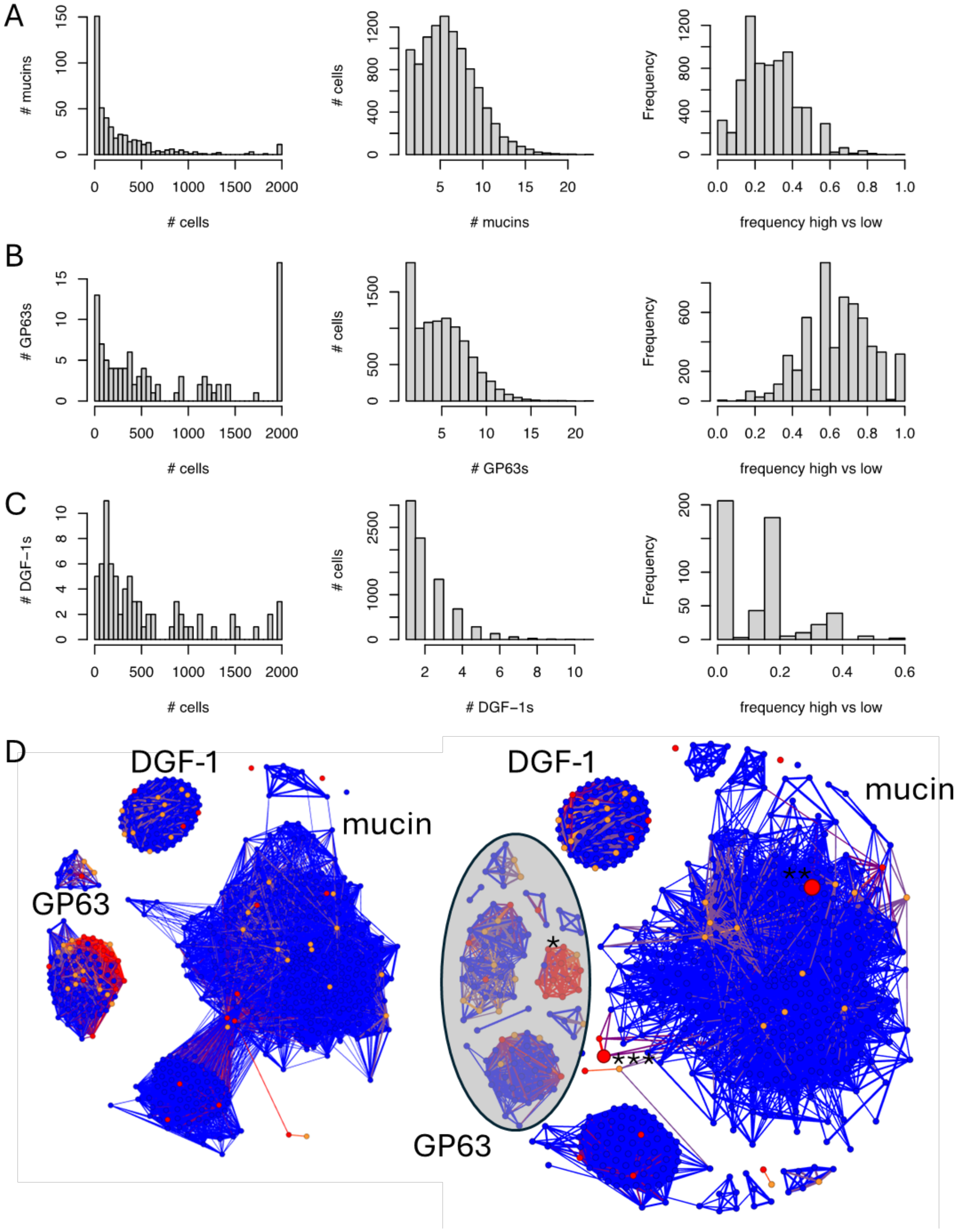

Expression pattern of 458 mucins (A), 96 Surface protease GP63 (B) and 75 dispersed gene family protein DGF-1 (C), see Supplementary Figure 3 for description. (D) Gephi graph for the three gene families with an identity cutoff of >21% (left) and >41% (right), as described in Supplementary Figure 3. At 21% identity, the three families cluster well together. At 41% the GP63 fall into several classes, with members expressed in more cells clustered together(*), as described for the other gene families before. However, for the mucins, C4B63_1g346 (**, large red dot) is expressed in 19,435 cells. C4B63_51g234 (second large dot ***), is expressed in 9,451 cells. They do not form a separate cluster of "conserved/highly expression" genes, compared to the other gene families. The DGF-1 genes are more conserved and share >80% sequence similarity.

**Supplementary Figure 6**. rRNA mapping

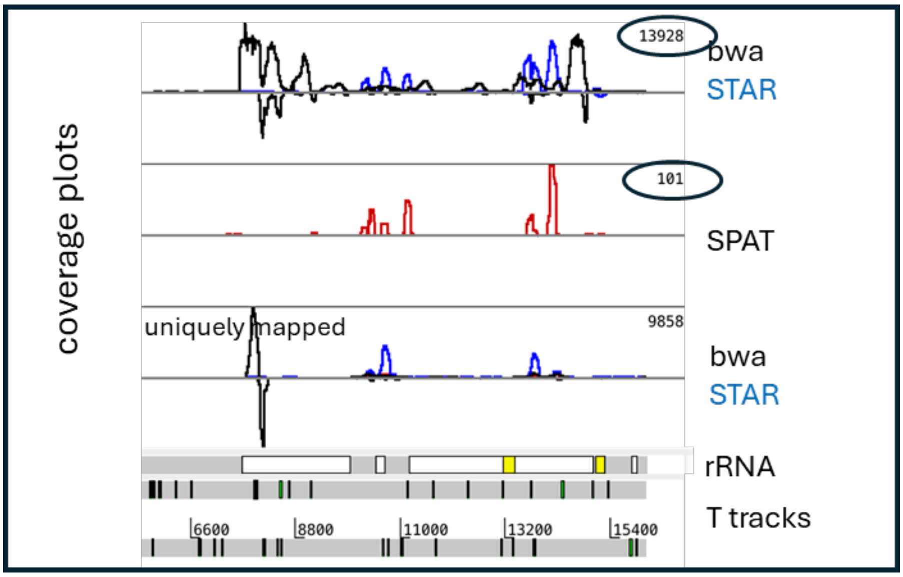


The white/yellow boxes are rRNA. The two plots how that the STAR mapper (blue lines), as part of cellRanger, maps far less reads compared to bwa (black line). There is no evidence of poly-adenylation of rRNA, as there are very few SPAT reads (<1%, noise). Rather, we hypothesize that due to the high abundance of rRNA in the parasite, small poly T/A tracks >5 bp (vertical lines, bottom of figure) are sufficient to anneal the rRNA to the capture oligos in the 10X reaction.

**Supplementary Table 1**. Sample information for scRNA-seq samples

| Sample Id | Stage collected | Viability (%) | No. cells/stage | No. cells/sample[c] | cDNA (ng/ul)[d] | library (ng/ul)[e] | Sequencing depth (Gb) |
|---|---|---|---|---|---|---|---|
| AMA6/24 | AMA6h | 91.6 | 8250 | 16500 | 2.2 | 8.1 | 220 Gb |
| | AMA24h | 87.3 | 8250 | | | | |
| EP/AMA120 | EP | 98.9 | 8250 | 16500 | 2.7 | 14.5 | 220 Gb |
| | AMA120h | 99.6 | 8250 | | | | |
| ADT | ADT | 99.5 | 16500 | 16500 | 2.9 | 17.3 | 220 Gb |
| SE/MT[a] | SE/MT | 97.8 | 16500 | 16500 | 0.8 | 7.8 | 70 Gb |
| | | 99.5 | 16500 | 16500 | 0.7 | 8.5 | 70 Gb |
| MIX[b] | AMA6h | 96.4 | 3500 | 21000 | 0.8 | 24.8 | 200 Gb |
| | AMA24h | 97.4 | 3500 | | | | |
| | AMA120h | 98.9 | 3500 | | | | |
| | EP | 98.2 | 3500 | 21000 | 0.7 | 29.2 | 200 Gb |
| | ADT | 98.9 | 3500 | | | | |
| | SE/MT | 96.7 | 3500 | | | | |

[a] Two biological replicates collected
[b] Two technical replicates collected
[c] Number of cells loaded into the 10X Chromium Next Gem Chip G
[d] Diluted in 40 µl total volume
[e] Libraries prepared with 10 µl cDNA; diluted in 35 µl total volume

**Supplementary Table 2**. Comparison of mapping stats for Dm28c 2018 and Sylvio X10 2017 for the MIX 1 dataset

| Reference genome[a] | Median genes per cell | No. cells | % reads mapping to the genome | % reads mapping confidently to transcriptome |
|---|---|---|---|---|
| Dm28c 2018 | 216 | 5464 | 55.0 | 4.7 |
| Sylvio X10 2017 | 135 | 5804 | 15.8 | 2.0 |

[a] The genomes have not been altered in any way from when they were downloaded from TriTrypDB (i.e. no UTR extension).

**Supplementary Table 3**. Selected statistics generated by 10X Cellranger output for scRNA-seq samples

| **Sample Id** | **Total reads** | **Total reads *T. cruzi*[a]** | **Median genes *T. cruzi*** | **% reads mapping confidently to:** | | | | |
|---|---|---|---|---|---|---|---|---|
| | | | | ***T. cruzi* transc (raw)** | ***T. cruzi* transc (adjusted)** | **human transc** | ***T. cruzi* genome** | **human genome** |
| AMA6/24 | 952950562 | 387850879 | 307 | 1.6 | 3.9 | 33.1 | 6.3 | 59.3 |
| EP/AMA120 | 682978286 | 678197438 | 928 | 14.4 | 14.5 | 0.2 | 43.1 | 0.7 |
| ADT | 729447233 | 729447233 | 762 | 7.7 | 7.7 | NA | 30.1 | NA |
| SE/MT 1 | 271060069 | 271060069 | 298 | 18.4 | 18.4 | NA | 44.4 | NA |
| SE/MT 2 | 251411209 | 251411209 | 421 | 21.4 | 21.4 | NA | 57.4 | NA |
| MIX 1 | 674582011 | 611171302 | 503 | 10.1 | 11.2 | 6.6 | 56.2 | 9.4 |
| MIX 2 | 739240858 | 663838290 | 422 | 9.9 | 11.0 | 7.3 | 56.8 | 10.2 |

[a] For datasets which contain human cells, 'total reads *T. cruzi*' was calculated by assuming all human reads will map to the human genome and the rest of the reads will be from the *T. cruzi* samples. This was then used to calculate the '% of reads mapping confidently to *T. cruzi* transc adjusted' values. Transc: transcriptome.

**Supplementary Table 4**. Cutoffs used for filtering out low quality cells across the scRNA-seq samples

| Sample Id | Gene counts minimum | Gene counts maximum | % reads mapping to *T. cruzi* genes maximum | % reads mapping to human transcriptome maximum |
|---|---|---|---|---|
| AMA6/24 | 200 | 750 | 20 | 50 |
| EP/AMA120 | 400 | 2000 | 30 | 10 |
| ADT | 200 | 1500 | 25 | NA |
| SE/MT 1 | 150 | 1000 | 40 | NA |
| SE/MT 2 | 200 | 1250 | 40 | NA |
| MIX 1 | 150 | 1500 | 28 | 30 |
| MIX 2 | 200 | 1250 | 30 | 30 |

**Supplementary Table 5**. Weighted average F1-scores for the scPoli predictions across the 10 atlas clusters, for the different subsets

| Cluster Id | 0.05_1[a] | 0.05_2 | 0.25_1 | 0.25_2 | 0.5_1 | 0.5_2 | 0.75_1 | 0.75_2 | 0.9_1 | 0.9_2 |
|---|---|---|---|---|---|---|---|---|---|---|
| early_mid_amast_5 | 0.80 | 0.74 | 0.73 | 0.73 | 0.71 | 0.73 | 0.63 | 0.62 | 0.43 | 0.47 |
| epi_meta_trans_6 | 0.75 | 0.77 | 0.76 | 0.75 | 0.75 | 0.76 | 0.70 | 0.65 | 0.47 | 0.44 |
| epi_meta_trans_7 | 0.59 | 0.61 | 0.59 | 0.70 | 0.60 | 0.63 | 0.54 | 0.57 | 0.39 | 0.31 |
| epimast_2 | 0.87 | 0.85 | 0.86 | 0.85 | 0.83 | 0.84 | 0.82 | 0.81 | 0.77 | 0.79 |
| late_amast_1 | 0.80 | 0.77 | 0.76 | 0.79 | 0.73 | 0.74 | 0.69 | 0.66 | 0.57 | 0.56 |
| meta_3 | 0.85 | 0.82 | 0.79 | 0.78 | 0.79 | 0.78 | 0.76 | 0.74 | 0.73 | 0.75 |
| meta_8 | 0.79 | 0.75 | 0.79 | 0.79 | 0.74 | 0.66 | 0.63 | 0.66 | 0.60 | 0.61 |
| trypomast_0 | 0.76 | 0.78 | 0.75 | 0.75 | 0.75 | 0.75 | 0.73 | 0.73 | 0.72 | 0.71 |
| trypomast_4 | 0.72 | 0.75 | 0.72 | 0.73 | 0.70 | 0.70 | 0.68 | 0.68 | 0.67 | 0.65 |
| trypomast_9 | 0.64 | 0.70 | 0.68 | 0.69 | 0.73 | 0.41 | 0.63 | 0.60 | 0.41 | 0.29 |

[a] The number before the underscore represents the proportion of the atlas that was used as the testing data. The number after the underscore indicates the sampling number.

**Supplementary Table 6**. Sample information for bulk RNA-seq samples

| Sample Id | No. parasites collected[a] | RNA (ng/µl[b]) | RNA (ng/1E+06 parasites) | mean RNA/stage (ng/1E+06 parasites) |
|---|---|---|---|---|
| bAMA6 | 8.40E+06 | 56 | 200 | 185.8 |
| | 7.52E+06 | 43 | 171.5 | |
| bAMA24 | 3.76E+06 | 22 | 175.5 | 146.9 |
| | 3.81E+06 | 15 | 118.3 | |
| bAMA120 | 1.16E+07 | 70 | 181 | 163.2 |
| | 1.94E+07 | 94 | 145.4 | |
| bEP | 4.00E+07 | 265 | 198.8 | 213.8 |
| | 4.00E+07 | 305 | 228.8 | |
| bADT | 4.00E+07 | 89 | 66.8 | 65.8 |
| | 4.00E+07 | 72 | 54 | |
| | 4.00E+07 | 102 | 76.5 | |
| bSE/MT | 4.00E+07 | 59 | 44.3 | 39 |
| | 4.00E+07 | 45 | 33.8 | |

[a] Two biological replicates collected per sample, with two technical replicates prepared for the second bADT biological replicate

[b] Diluted in 30 µl total volume

**Supplementary Table 7**. Mapping statistics of the bulk RNA sequencing samples mapped against the Dm28c *T. cruzi* + kDNA genome

| Sample Id | No. input reads[a] | % uniquely mapped reads | % reads mapped to multiple loci | % reads mapped to too many loci | % reads unmapped: too many mismatches | % reads unmapped: too short | % reads unmapped: other | Total alignments (featureCounts) | % successfully assigned alignments (featureCounts) |
|---|---|---|---|---|---|---|---|---|---|
| bAMA6 | 23475929 | 41.88 | 14.48 | 9.83 | 0 | 31.79 | 2.02 | 21942505 | 38.20 |
| | 20219803 | 46.33 | 16.26 | 10.92 | 0 | 24.33 | 2.17 | 21107762 | 38.20 |
| bAMA24 | 20498527 | 36.14 | 12.46 | 5.11 | 0 | 44.57 | 1.73 | 16395372 | 38.10 |
| | 24492333 | 35.19 | 12.00 | 4.52 | 0 | 46.42 | 1.87 | 18848721 | 39.50 |
| bAMA120 | 26077005 | 52.40 | 17.93 | 16.22 | 0 | 9.89 | 3.56 | 30285994 | 36.40 |
| | 27749110 | 49.86 | 17.13 | 21.31 | 0 | 8.70 | 2.99 | 31157951 | 35.00 |
| bEP | 25717422 | 54.56 | 19.39 | 18.14 | 0 | 4.01 | 3.90 | 32736362 | 34.30 |
| | 25985438 | 56.38 | 18.93 | 16.58 | 0 | 3.91 | 4.20 | 32444213 | 36.80 |
| bADT | 27090531 | 51.03 | 20.71 | 22.63 | 0 | 3.91 | 1.73 | 34960443 | 31.40 |
| | 21882358 | 48.82 | 20.93 | 22.23 | 0 | 5.18 | 2.84 | 27812588 | 31.80 |
| | 22048702 | 43.12 | 19.93 | 28.85 | 0 | 5.46 | 2.64 | 26768688 | 29.30 |
| bSE/MT | 22249655 | 59.79 | 18.83 | 14.13 | 0 | 4.02 | 3.23 | 28940501 | 36.50 |
| | 28225858 | 64.08 | 18.66 | 9.96 | 0 | 4.78 | 2.52 | 36415986 | 39.50 |

[a] Two biological replicates were collected per sample, with two technical replicates prepared for the second bADT biological replicate
